# Bidirectional and reciprocal control of astrocyte free calcium by modest fluctuations in external potassium

**DOI:** 10.1101/2022.08.08.503209

**Authors:** A. Institoris, S. Shin, N.L. Weilinger, K. Gorzo, E. Mehina, J. Haidey, B.A. MacVicar, G.R. Gordon

**Affiliations:** Hotchkiss Brain Institute, Department of Physiology and Pharmacology, Cumming School of Medicine, University of Calgary, AB, Calgary, Canada

**Author notes:** Corresponding Author: Grant R. Gordon, Room 1B40A, Building HRIC, 3330 Hospital Dr. NW, Calgary, AB, Canada, T2N 4N1. authors contributed equivalently (A. Institoris, S. Shin).

**Keywords:** astrocyte, calcium, potassium, two-photon, FLIM, brain slice, in vivo, arteriole

## Abstract

Astrocytes sense and respond to changes in the concentration of extracellular K^+^, and separately contribute to multiple physiological processes through Ca^2+^ dependent mechanisms. Yet, whether a modest change in [K^+^]_o_ impacts astrocyte free Ca^2+^ remains unclear. Using relative or quantitative two-photon fluorescence Ca^2+^ imaging in acute brain slices or *in vivo* in the somatosensory cortex from Sprague Dawley rats and C57Bl/6 mice, we showed that changes to external K^+^ (+/-1mM to 2.5mM) reciprocally controls the astrocyte Rhod-2 or OGB-1 Ca^2+^-dependent fluorescence in the soma, major processes and endfeet. The astrocyte Ca^2+^ decrease when [K^+^]_o_ was elevated was sensitive to lowering the external concentration of Ca^2+^, Cl^-^, and HCO_3_^-^, but not Na^+^. Unexpectedly, the phenomenon was blocked by inhibiting K-Cl cotransport. Picrotoxin induced ictal neural activity drove an analogous decrease of astrocyte Ca^2+^. K^+^ mediated cerebral arteriole dilation in brain slices was also sensitive to inhibiting K-Cl cotransport as well as whole-cell patching a peri-arteriole astrocyte which perturbs normal Ca^2+^, Cl^-^ and HCO_3_^-^ concentration gradients. These data reveal subtle, bidirectional regulation of astrocyte free Ca^2^ via fluctuations of [K^+^]_o_ within the physiological range.

## Introduction

Astrocytes are important sensors and regulators of the extracellular milieu, including potassium (K^+^) homeostasis. How changes in the external K^+^ concentration impact brain function have been investigated for over 80 years. Early measurements using K^+^-selective microelectrodes showed that external K^+^ fluctuates around 3mM *in vivo* (Somjen, 1979) and that physiological neuronal action potential firing raises [K^+^]_o_ typically from baseline by 0.1-1mM (Kelly and Van Essen, 1974; Syková et al., 1974; Singer and Lux, 1975; Korytová, 1977). Similar sized shifts in baseline extracellular K^+^ are observed switching between sleep and wake states (Ding et al., 2016), but higher elevations likely occur in microenvironments. Postsynaptic NMDA receptor opening elevates extracellular K^+^ with elevations modelled to reach between 5-7 mM for tens of milliseconds in the synaptic cleft (Shih et al., 2013). Astrocytes help take up these [K^+^]_o_ increases via Na^+^/K^+^ ATPase and K_ir_ 4.1 channel activity (Hertz et al., 2015; Chever et al., 2010) to help maintain neuronal ionic gradients (Kofuji and Newman, 2004)(Hertz et al., 2015)(Chever et al., 2010). Astrocytes also depolarize to elevated [K^+^]_o_ which drives HCO_3_^-^ influx via the electrogenic Na^+^/HCO_3_^-^ cotransporter. This increases intracellular pH (pH_i_) (Ransom et al., 2000; Larsen and MacAulay, 2017) and causes downstream activation of soluble adenylyl cyclase (Choi et al., 2012). However, changes within or just beyond physiological levels of [K^+^]_o_ are not thought to affect astrocyte Ca^2+^ signaling, until pathological levels of [K^+^]_o_ (∼20mM) are reached (Duffy and MacVicar, 1994).

Indeed, astrocytes regulate an array of physiological functions through changes in free cytosolic Ca^2+^, including synaptic strength (Fiacco and McCarthy, 2004; Henneberger et al., 2010) and local cerebral blood flow (Mulligan and MacVicar, 2004; Takano et al., 2006; Rosenegger et al., 2015; Mishra et al., 2016; Haidey et al., 2021). While most studies have focused on large amplitude, transmitter evoked Ca^2+^ transients in astrocytes, a multitude of different types of Ca^2+^ signals likely exist in these cells (Srinivasan et al., 2015; Agarwal et al., 2017; Haidey et al., 2021) that are incompletely understood. For example, changes to the resting, or steady-state free [Ca^2+^]_i_ concentration in astrocytes is seldom explored. This is important because astrocytes have a relatively high resting free Ca^2+^ concentration in the soma and major processes (Zheng et al., 2015) and relatively small deviations from resting Ca^2+^ can impact gliotransmission (Parpura and Haydon, 2000; Shigetomi et al., 2012). Resting astrocyte Ca^2+^ can be modulated by changes in plasma membrane Ca^2+^ flux (Agarwal et al., 2017; Rungta et al., 2016; Shigetomi et al., 2011), arteriole tone (Haidey et al., 2021; Kim et al., 2015) or dopamine (Jennings et al., 2017). Additionally, bursts of afferent activity decrease steady-state astrocyte Ca^2+^ in an NMDA receptor-dependent manner (Mehina et al., 2017). As NMDA receptor opening (Shih et al., 2013) or neuromodulators (Ding et al., 2016) can affect the external potassium concentration, we tested the hypothesis modest changes in [K^+^]_o_ regulate the resting [Ca^2+^]_i_ in astrocytes. This exploration is needed because a small shift in astrocyte free Ca^2+^ may have been previously overlooked using standard fluorescence techniques to measure Ca^2+^ that did not account for the changes to astrocyte volume that accompany changes in [K^+^]_o_ (Florence et al., 2012).

## Results

### Modest changes in [K^+^]_o_ reciprocally shifts free [Ca^2+^]_i_ in astrocytes

Using two-photon fluorescence imaging, we bath applied isosmotic high [K^+^]_o_ solutions onto acute slices of the somatosensory cortex from Sprague Dawley rats. Slices were bulk loaded with the bright, relatively high-affinity Ca^2+^ indicator Rhod-2/AM, along with the morphological dye Calcein/AM. We took the ratio of the normalized Rhod-2 fluorescence signal over the normalized Calcein fluorescence signal to correct for apparent decreases in Ca^2+^ that were the result of cell swelling (Florence et al., 2012) (Figure 1A). Indeed, we observed decreases in both the Rhod-2 Ca^2+^ signal and Calcein signal in response to a +2.5mM [K^+^]_o_ challenge (going from 2.5mM to 5mM K+), yet interestingly, the drop in Rhod-2 fluorescence was nearly double that of the morphological fluorescence (Figure 1B). This suggested a decrease in [Ca^2+^]_i_ beyond that predicted by a cell volume increase alone. The Rhod-2/Calcein ratio revealed consistent and significant decreases in astrocyte steady-state Ca^2+^ when [K^+^]_o_ was increased by +2.5mM to 5 mM (0.83 ± 0.1, n=9 slices, p=0.001, Figure 1C-E). Even smaller challenges of only +1mM (increasing [K^+^]_o_ from 2.5mM to 3.5mM) decreased Ca^2+^ (0.88 ± 0.01, n=4, p=0.002, Figure 1H) but to a lesser degree than a +2.5mM challenge. Moving beyond the physiological range, a +5mM challenge (2.5 to 7.5mM) caused a proportionally larger drop in astrocyte Ca^2+^ (0.66 ± 0.02, n=4, p=0.002, Figure 1H). Observations of up and down shifts in [K^+^]_o_ *in vivo* (Ding et al., 2016), let us to test whether decreasing [K^+^]_o_ would produce an opposite effect. For this experiment, we maintained our slices in 3.5mM K^+^ and decreased [K^+^]_o_ to 2.5mM (−1mM K^+^ challenge) and found a significant increase in astrocyte free Ca^2+^ (1.09 ± 0.02, n=5, p=0.007, Figure 1F,G)(dose response summary Figure 1H). These data suggest that external [K^+^]_o_ controls the resting free Ca^2+^ concentration in astrocytes in a bidirectional manner.

**Figure 1:**
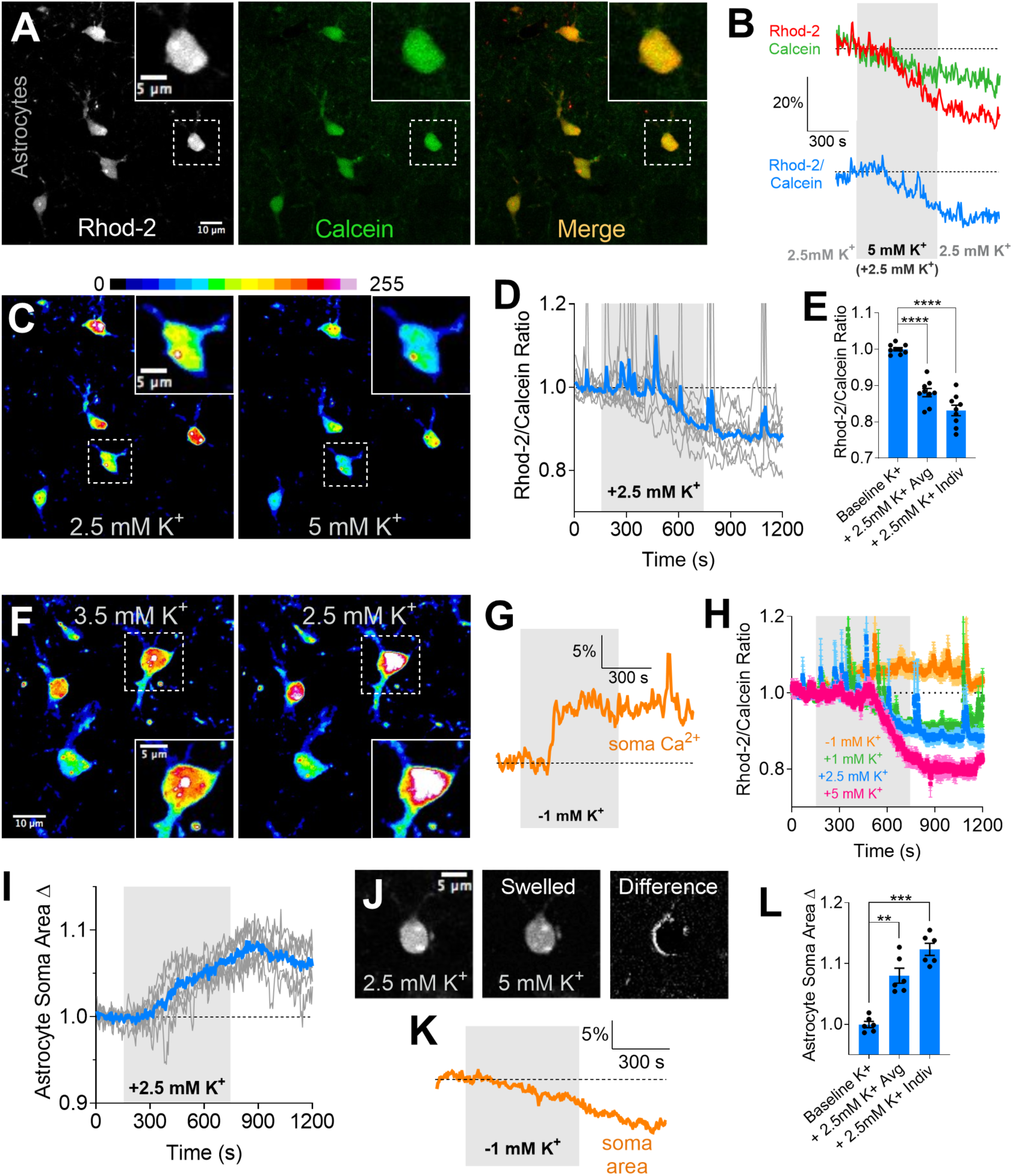
Modest changes in external [K^+^] bidirectionally control astrocyte free Ca^2+^ and soma volume. **A**) Astrocytes in an acute somatosensory cortical brain slice co-loaded with Rhod-2/AM (left) and Calcein/AM (middle), merge (right). Inset images show a close-up of a single astrocyte soma. **B**) Normalized Rhod-2 and Calcein traces (upper) measured from an astrocyte soma and the corresponding ratio (lower) in response to a modest [K^+^]_o_ challenge (+2.5mM): from 2.5mM to 5mM (grey bar) back to 2.5mM. **C**) Pseudo coloured Rhod-2 images showing the decrease in Ca^2+^ signal to a +2.5mM [K^+^]_o_ increase. **D**) Ca^2+^ trace summary showing individual experiment traces (grey) and overall average (blue, without error). **E**) Ca^2+^ summary showing the average peak effect (same time point for all) and the peak of effect of individual experiments (peaks at different time points for each) for the +2.5mM [K^+^]_o_ increase. **F**) Pseudo coloured Rhod-2 images showing the increase in astrocyte Ca^2+^ signal to a −1.0mM [K^+^]_o_ decrease. **G**) Ca^2+^ trace from a single experiment showing the Ca^2+^ increase from a [K^+^]_o_ decrease of 1mM. **H**) Ca^2+^ summary trace data of the dose response to [K^+^]_o_ changes, showing bidirectional effects. **I**) Astrocyte soma area measures showing individual experiment traces (grey) and overall average (blue, without error) to the +2.5mM [K^+^]_o_ challenge. **J**) Images depicting the soma volume increase. Difference image subtracts the large volume astrocyte from the small volume state. **K**) Soma area trace from one experiment showing a decrease in cell area when [K^+^]_o_ is decreased by 1mM (3.5 to 2.5mM). **L**) Soma area summary showing the average peak effect (same time point) and the peak of effect of individual experiments (different time point for each) for the +2.5mM [K^+^]_o_ increase experiments. Data is mean +/-SEM, paired two-tailed, t-tests. ** p<0.01, *** p<0.001

We confirmed a previously described volume increase in astrocytes to a modest elevation in [K^+^]_o_. A +2.5mM [K^+^]_o_ challenge increased soma area (1.12 ± 0.01, p=0.001, n=6, Figure 1I,J, L), which also supported our observed decrease in the morphological Calcein signal. However, expanding on previous findings, we found these volume changes were bidirectional in nature and occurred in response to smaller changes in [K^+^]_o_, similar to what we observed for astrocyte Ca^2+^. For example, a −1mM [K^+^]_o_ test, decreased soma area (Figure 1K). Collectively, these data show that [K^+^]_o_ within and slightly beyond the physiological range bidirectionally controls free Ca^2+^ in astrocytes, more so than expected from volume changes alone.

### FLIM reveals [K^+^]_o_-mediated decrease in astrocyte free [Ca^2+^]_i_

To test the idea that a modest increase in external K^+^ causes a bon-a-fide decrease in the free Ca^2+^ concentration in astrocytes, we performed two-photon Fluorescence Lifetime Imaging Microscopy (FLIM)(Figure 1A). This is a quantitative method for assessing cell [Ca^2+^]_i_ in which measurements are independent of dynamic changes in dye concentration, photobleaching or to changes in optical properties of tissue. We used the high affinity, FLIM-sensitive Ca^2+^ indicator OGB-1 to circumvent potential problems associated with relative two-photon fluorescence intensity measurements using Rhod-2 and Calcein during K^+^-induced astrocyte swelling. Our calibration of the OGB-1 FLIM decay to various levels of free [Ca^2+^] in solution (see methods) placed the Kd at 192 nM (Figure 2C), ideal to detect small shifts in resting astrocyte free [Ca^2+^]. We patch loaded single astrocytes with OGB-1, which readily diffused to neighboring astrocytes that were coupled via gap junctions to the patched cell. We allowed 10 min for OGB-1 diffusion through the astrocyte network. As previously reported (Zheng et al., 2015), the patched astrocyte displayed elevated resting [Ca^2+^] compared to the adjacent gap junction (GJ) coupled astrocytes (patched soma: 115.6 ± 9.7 nM; GJ soma: 69.5 ± 5.5 nM, n=8, p=0.003, Figure 2B,D,E). Similar to our results using the Rhod-2/Calcein ratio, bath application of an isosmotic +2.5mM K^+^ challenge (2.5 to 5mM K+) decreased the OGB-1 FLIM Ca^2+^ signal (shortened the mean lifetime) in the soma and major processes of both the patched astrocyte and coupled astrocytes (normalized GJ somata: 0.72 ± 0.05, n=8 p=0.001; Figure 2B,D,E). We also noted a decreased resting [Ca^2+^]_i_ level in astrocytic perivascular endfeet (normalized: 0.71 ± 0.04, p=0.008, n=4, Figure 2F,G), in response to high [K^+^]_o_. We then examined astrocytic endfeet apposed to blood vessels using our Rhod-2/Calcein approach and observed a similar decrease in Ca^2+^ caused by +2.5mM high [K^+^]_o_ (0.74 ± 0.01, n=3, Figure 2H,I) and clear endfoot swelling (Figure 2J). These data suggest that a modest elevation in external K^+^ causes a genuine decrease in the free Ca^2+^ concentration in astrocytes in multiple large cellular compartments.

**Figure 2:**
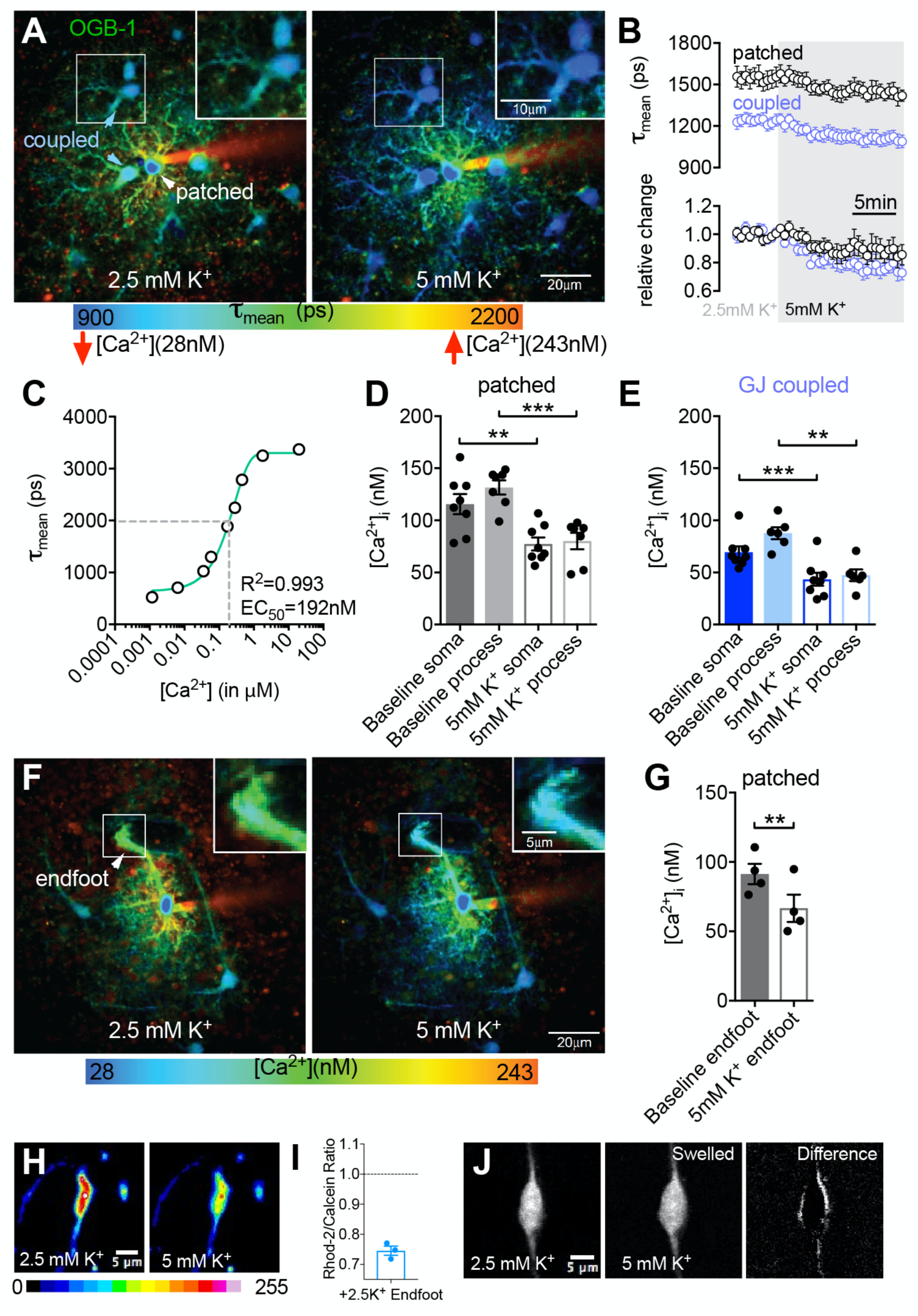
Fluorescence Lifetime Imaging Microscopy (FLIM) reveals a quantitative decrease in astrocyte free Ca^2+^ in response to a modest elevation in external K^+^. **A**) Two-photon FLIM image sequence showing a patched astrocyte loaded with OGB-1, and coupled astrocyte also loaded, in response to a +2.5mM K+ increase. Pseudo colouring corresponds to the tau mean of the lifetime decay curves (cooler colours: faster decay = lower Ca^2+^; warmer colours: slower decay = higher Ca^2+^). High K^+^ results in a shift towards faster lifetimes (cooler colours) thus decreased free Ca^2+^ in both the patched astrocyte and the coupled astrocytes. **B**) (Upper) Raw, averaged OGB-1 lifetimes from patched astrocytes (black) and coupled astrocytes (blue) over time in response to high K^+^. (Lower) Normalized lifetime values over time for the same data. **C**) OGB-1 Lifetime-Ca^2+^ concentration calibration curve. **D**) Summary of Ca^2+^ concentrations in patched astrocyte somata and major processes before and during high K^+^. **E**) Summary of Ca^2+^ concentrations in coupled astrocyte somata and major processes before and during high K^+^. **F**) FLIM images of a patched, peri-vascular astrocyte before and during high K^+^, inset shows endfoot. **G**) Summary of Ca^2+^ concentrations in patch loaded endfeet before and during high K^+^. **H**) Rhod-2 images showing the drop in astrocyte Ca^2+^ occurs in perivascular endfeet. **I**) Rhod2/Calcein ratio endfoot summary data. **J**) Images show an endfoot swelling in response to high K^+^. Data is mean +/-SEM, paired two-tailed t-tests, ** p<0.01, *** p<0.001

### Elevated [K^+^]_o_ decreases astrocyte Ca^2+^ *in vivo*

We sought to confirm whether the decrease in astrocyte free Ca^2+^ to high [K^+^]_o_ occurred under realistic conditions *in vivo*. We used an awake, but lightly sedated mouse model in which mice were head-fixed under the two-photon microscope (Bonder and McCarthy, 2014). Using chlorprothixene (2mg/kg), animals remain calm but awake during imaging. Furthermore, by using perforated cranial windows (Tran et al., 2018), we could superfuse isotonic high K^+^ solutions onto the surface of neocortex through a ∼600 micron diameter hole, while imaging within the hole itself. Though a +/-1mM K^+^ change could be detected in acute slices, there is a well appreciated diffusion barrier for superfusion experiments on the neocortical surface crossing the pia mater, even with dura removal. Therefore, we attempted to measure changes in the astrocyte Ca^2+^ level in response to a 10mM K^+^ solution. Indeed, elevating K^+^ in this manner again resulted in a prominent decrease in Rhod-2 fluorescence in astrocyte somas and other major compartments that were within the coverglass hole in the superficial neocortex (0.69 ± 0.03, N=3, p =0.006, Figure 3). Though these experiments were not controlled for by Calcein co-loading, we observed little astrocyte swelling *in vivo* (1.01 ± 0.03, N=3). These data demonstrate a similar astrocyte phenomenon *in vivo*, to our effects reported in acute brain slices. They also suggest there is a drop in free Ca^2+^ independent of dilution from swelling.

**Figure 3:**
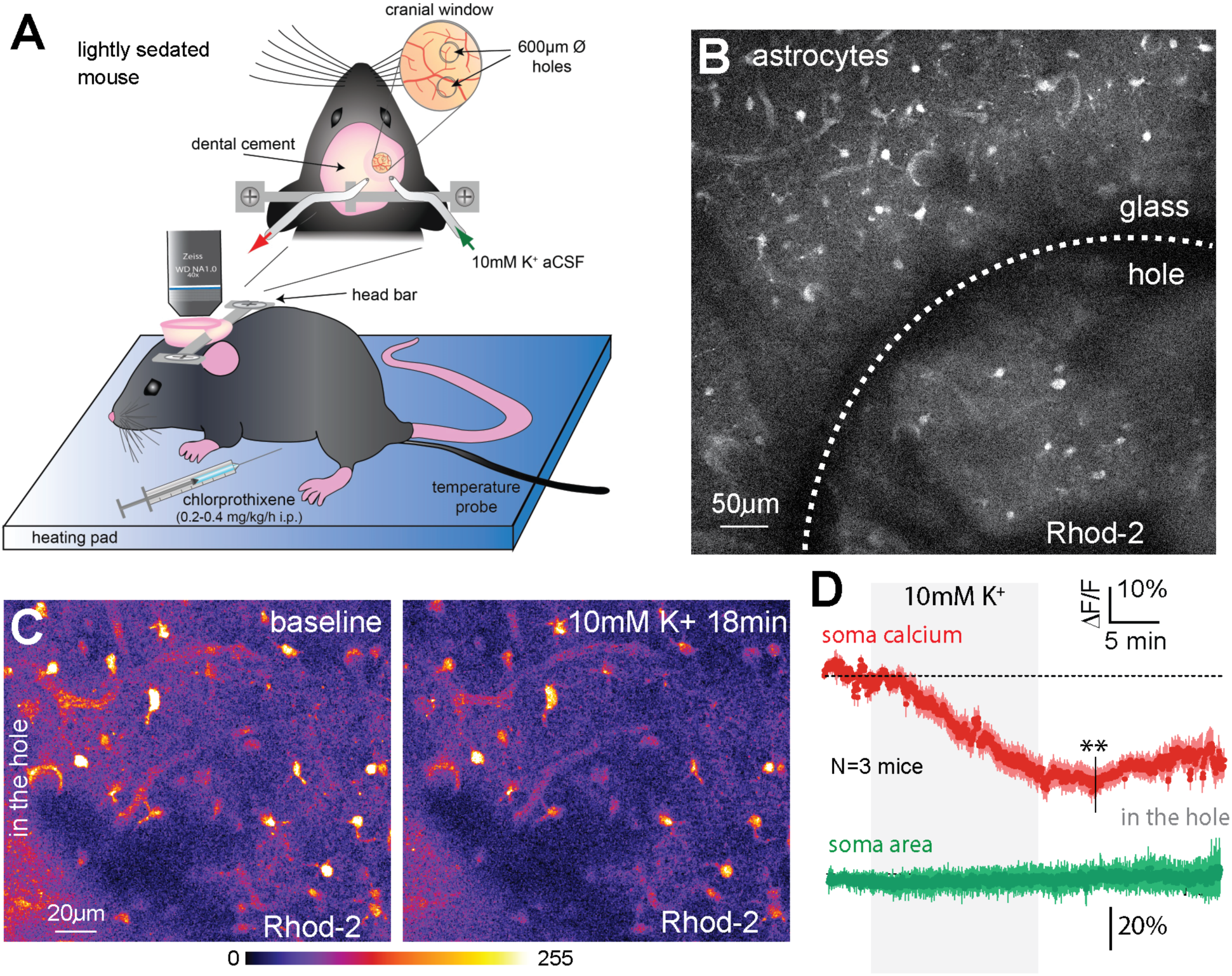
Elevated [K^+^]_o_ decreases astrocyte Ca^2+^ *in vivo*. **A**) Schematic of lightly sedated mouse *in vivo* setup. High K^+^ solution is superfused onto the surface of the brain via a perforated cranial window. **B**) Large field of view two-photon image of Rhod-2/AM loaded astrocytes, showing astrocytes located either within the coverglass hole or underneath the coverglass. **C**) Images of before and during treatment of 10mM K^+^. LUT coloured astrocytes show a decrease in Rhod-2 signal during high K^+^. **D**) Averaged trace data showing the decrease in astrocyte Rhod-2 signal in 14 astrocytes across 3 different mice in response to 10mM K^+^. In contrast, astrocytes soma area did not increase. Measurements were taken from astrocytes located within the coverglass hole. Data is mean +/-SEM, paired two-tailed t-tests, ** p<0.01.

### The K^+^-induced change to astrocyte free Ca^2+^ is not plastic

We extended the time frame of our K^+^ experiment and examined the washout period using Rhod-2 and Calcein. We observed a complete washout of the Ca^2+^ drop to a +2.5mM [K^+^]_o_ challenge (Supplementary Figure 1A). As a shift in [K^+^]_o_ of this magnitude has little impact on neural excitability (Somjen, 1979), we confirmed that the K^+^-induced drop in astrocyte Ca^2+^ was insensitive to TTX (500nM) (Supplementary Figure 1B). Theta-burst afferent synaptic activity also decreases astrocyte steady-state Ca^2+^, and this effect is occluded in enriched animals (Mehina et al., 2017). With a complete washout of the K^+^ effect and no reliance on neural action potential firing, we tested whether the K^+^-induced decrease in astrocyte Ca^2+^ would be unaltered after enrichment. Five Sprague Dawley rats were housed in an enrichment environment for three weeks (see methods) before acute slices were prepared and the same +2.5mM [K^+^]_o_ experiment was conducted. We found that enrichment had no effect (p=0.24) on the magnitude of the astrocyte Ca^2+^ decrease to high K^+^ when compared to our control data set (Supplementary Figure 1C-F). These data suggest that the astrocyte free Ca^2+^ decrease following a modest elevation in [K^+^]_o_ neither depends on changes in neural activity nor on long-lasting plastic mechanisms. Given the bidirectional nature of the effect described above, these data collectively suggest that changes to steady-state [K^+^]_o_ reciprocally controls steady-state free Ca^2+^ in astrocytes: when [K^+^]_o_ is elevated, astrocyte free Ca^2+^ decreases and when [K^+^]_o_ is decreased, astrocyte free Ca^2+^ increases.

### The decrease in astrocyte Ca^2+^ to moderate high K^+^ requires external Ca^2+^ and Bicarbonate

These observations directed us to explore whether external K^+^ affected the movement of Ca^2+^ across the astrocyte plasma membrane or the endoplasmic reticulum. It is well appreciated that increases in external K^+^ depolarize the astrocyte resting membrane potential which is predominately controlled by E_K_. Depolarizing membrane potential would decrease the driving force for Ca^2+^ entry, as well as alter the activity of electrogenic transporters. We tested whether the K^+^ effect relied on an external Ca^2+^ source or an internal one. Brain slices were preincubated with the potent SERCA pump inhibitor thapsigargin (1μM) for 30min to deplete intracellular Ca^2+^ stores. After this treatment we were unable to evoke large glutamate-mediated Ca^2+^ transients in astrocytes (Mehina et al., 2017). In this condition, a +2.5mM [K^+^]_o_ challenge still produced a Ca^2+^ decrease (p=0.21) measured by the Rhod-2/Calcein ratio (0.81 ± 0.01, n=11, Figure 4A,B), suggesting Ca^2+^ stores were not involved. In contrast, incubating our slices in a Ca^2+^ free ACSF external solution, significantly attenuated both the drop in free Ca^2+^ (0.90 ± 0.01, p=0.0015, n=8, Figure 4C,E,F) and astrocyte swelling (1.07 ± 0.01, p=0.001, n=7, Figure 4D,G,H) that normally occurred in response to high [K^+^]_o_.

**Figure 4:**
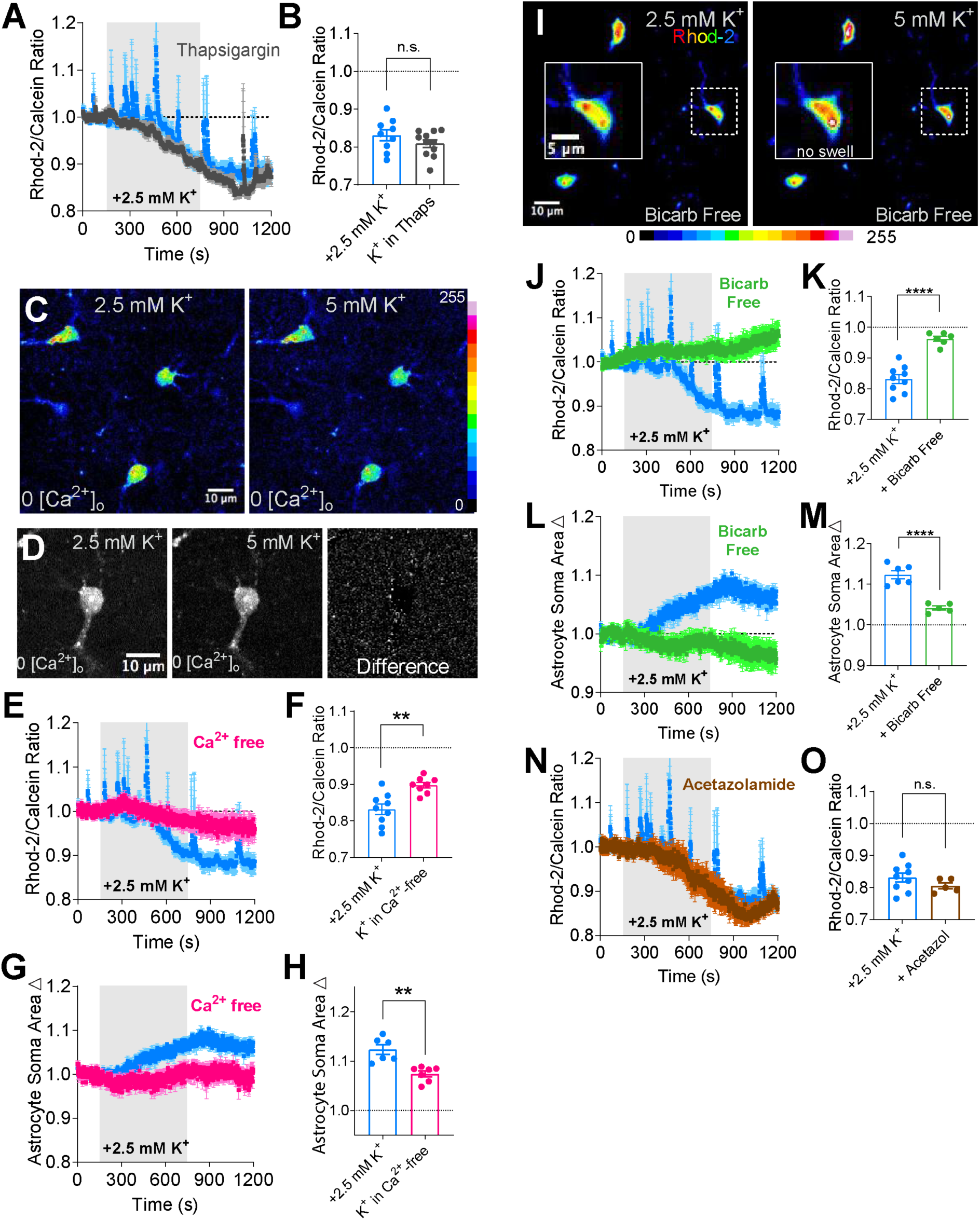
The [K^+^]_o_ effect on astrocytes depends external Ca^2+^ and Bicarbonate. **A**) Average summary time series Ca^2+^ data showing no effect on the K+-mediated decrease when internal Ca^2+^ stores are emptied with a 30min pretreatment with the SERCA pump inhibitor thapsigargin (1 μM). **B**) Summary of the peak decreases in Ca^2+^ from individual experiments. **C**) Psuedo coloured two-photon images of astrocytes loaded with Rhod-2 in a Ca^2+^ free external solution. A +2.5mM K^+^ challenge produced little change to resting astrocyte Ca^2+^. **D**) Image of an astrocyte at baseline K^+^ (left), elevated K^+^ (middle) and a difference image (5mM K^+^ minus 2.5mM K^+^) showing little change to astrocyte soma area in response to high K^+^ in zero external Ca^2+^. **E**) Average summary time series Ca^2+^ data in response to high K^+^ in a Ca^2+^ free external solution. **F**) Summary data of the maximal decrease in astrocyte Ca^2+^ in each experiment. **G**) Average summary time series soma area data in response to high K^+^ in a Ca^2+^ free external solution. **H**) Summary data of the maximal increase in astrocyte soma area in each experiment. **I)** Pseudo coloured two-photon images of astrocytes loaded with Rhod-2, showing no decrease in free Ca^2+^ to a +2.5mM K^+^ challenge when bicarbonate is removed from the external solution (HEPES buffered). Inset shows an astrocyte close up and the lack of cell swelling to high [K^+^]_o_ in a bicarb free external solution. **J**) Summary time series of Rhod-2/Calcein ratio Ca^2+^ data of the same experiment in *(I)* compared to control. **K**) Summary data of peak decreases in Ca^2+^ from each experiment. **L**) Summary time series soma area data in response to high [K^+^]_o_ in a bicarbonate free external solution. **M**) Summary data of peak increases in soma area from each experiment. **N**) Summary time series of Rhod-2/Calcein ratio Ca^2+^ data in response to high [K^+^]_o_ in the presence of acetazolamide (100 μM) compared to control. **O**) Summary data of peak decreases in Ca^2+^ in each experiment from acetazolamide vs control. Data is mean +/-SEM, unpaired two-tailed t-tests, ** p<0.01, **** p<0.0001

Bicarbonate movement into astrocytes plays a major role in volume regulation (Florence et al., 2012; Larsen and MacAulay, 2017) and pH alkalization (Ransom et al., 2000; Zhou et al., 2010; Larsen and MacAulay, 2017) during K^+^ challenges. To test whether bicarbonate affected astrocyte Ca^2+^, we substituted bicarbonate with HEPES in the ACSF. Notably, in bicarb-free conditions, a +2.5mM K^+^ challenge failed to decrease astrocyte free Ca^2+^ (0.96 ± 0.01, p=0.001, n=6, Figure 4I-K) and, as previously reported, blocked cell swelling (1.04 ± 0.01, p=0.001, n=5, Figure 4L,M). Cell swelling also depends on the pH change via the action of carbonic anhydrase (Florence et al., 2012) but a reliance on pH for Ca^2+^ changes is unclear. We found that blocking carbonic anhydrase activity with acetazolamide (100 μM) had no impact on the drop in astrocyte Ca^2+^ observed in response to high [K^+^]_o_ (0.80 ± 0.01, p=0.24, n=5, Figure 5N,O). These data suggest that bicarbonate, but not pH per se, plays an important role in the free Ca^2+^ decrease observed in response to a modest [K^+^]_o_ increase.

**Figure 5:**
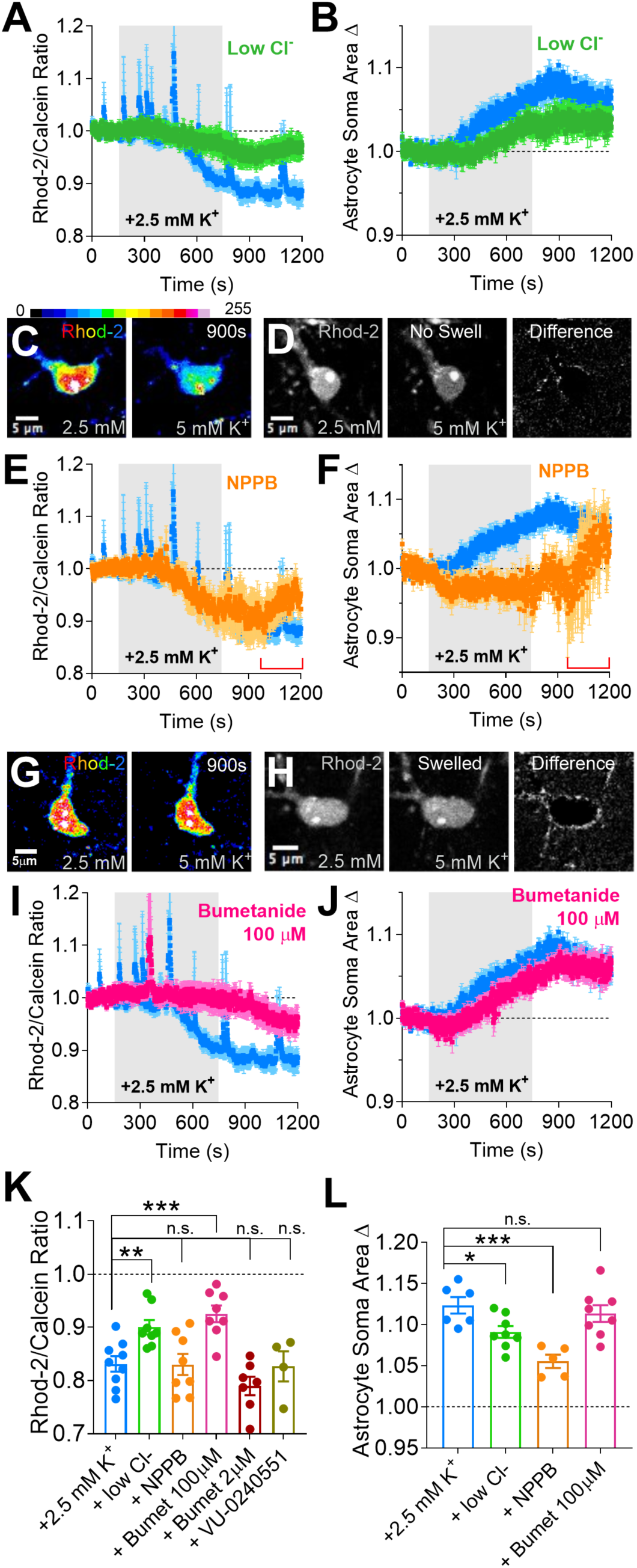
Anion channels and K-Cl co-transport separately control cell swelling and the Ca^2+^ decrease in response to moderate high [K^+^]_o_. **A)** Summary time series of Rhod-2/Calcein ratio Ca^2+^ data in response to high [K^+^]_o_ in the presence of a low Cl^-^ external solution (7.5 mM) compared to control. **B**) Summary time series of soma area in response to high [K^+^]_o_ in a low external Cl^-^ external solution. Both the Ca^2+^ drop and cell swelling were reduced in low Cl^-^ ACSF. **C**) Pseudo coloured two-photon images of an astrocyte loaded with Rhod-2, showing the decrease in free Ca^2+^ to a +2.5mM K^+^ challenge when anion channels were blocked with NPPB (100 μM). **D**) Two-photon images of an astrocyte showing the high K^+^-induced cell swelling was blocked in NPPB: no border or edge in difference image. **E and F**) Summary time series of Rhod-2/Calcein ratio Ca^2+^ data *(E)* and astrocyte soma area *(F)* in the same NPPB experiment depicted in *(C)* and *(D)*. Red brackets indicated period where there was tissue distortion occurring across the brain slice, compromising our measures. **G**) Pseudo coloured images of a Rhod-2 loaded astrocyte showing little decrease in free Ca^2+^ to a +2.5mM K^+^ challenge in the presence of the potassium chloride co-transporter blocker bumetanide (100 μM). **H**) Images of an astrocyte showing that the high K^+^ induced cell swelling still occurs in bumetanide. **I and J**) Summary time series data of astrocyte Ca^2+^ *(E)* and astrocyte soma area *(F)* in the same bumetanide experiment depicted in *(G)* and *(H)*. **K**) Summary data of peak decreases in astrocyte Ca^2+^ in each experiment. **L**) Summary data of peak increases in astrocyte soma area in each experiment. Data is mean +/-SEM, unpaired two-tailed t-tests, * p<0.05, ** p<0.01, *** p<0.001

### The change in astrocyte Ca^2+^ by [K^+^]_o_ does not rely on K_ir_ or SLCA4A

Astrocytes buffer the extracellular space from K^+^ increases, partly through inward rectifying potassium 4.1 channels (Chever et al., 2010). To test for the involvement of these channels in the K^+^-induced Ca^2+^ decrease, we tested high [K^+^]_o_ in the presence of BaCl_2_ (100 μM), which failed to block the effect (Supplementary Figure 2A). Next, we pondered whether K^+^-induced depolarization engaged the electrogenic negative sodium bicarbonate cotransporter SLCA4A (O’Connor et al., 1994) to activate soluble adenylyl cyclase (Choi et al., 2012). To explore this pathway, first we tested the +2.5mM K^+^ change in the presence of the SLCA4A antagonist S0589 (100μM) but found no attenuation in the decrease in astrocyte Ca^2+^ (Supplementary Figure 2B). Next we tested modest high [K^+^]_o_ in the presence of the soluble adenylyl cyclase inhibitor KH7 (30μM), but again found no effect on the drop in Ca^2+^ measured by the Rhod-2/Calcein ratio (Supplementary Figure 2C). To more broadly probe for the involvement of transporters or exchangers that rely on Na^+^ influx, GLT-1, GLAST, Na/K/Cl cotransporter or the Na^+^/Ca^2+^ exchanger, we tested high [K^+^]_o_ in the presence of a low Na^+^ external ACSF solution. By replacing NaCl with choline-chloride ([Na^+^]_o_ decreasing from 152.25mM to 26.25mM), we still failed to find a significant reduction in the decrease in free astrocyte Ca^2+^ caused by high [K^+^]_o_ (p=0.06, Supplementary Figure 2D), potentially ruling out channels and ion transport mechanisms that rely on the driving force of Na^+^ entry.

### Potassium and chloride cotransport is necessary for the drop in astrocyte Ca^2+^

K^+^ and Cl^-^ are moved simultaneously across the plasma membrane by the sodium potassium chloride co-transporter (NKCC) and the potassium chloride cotransporter (KCC). Other K^+^ transport mechanisms such as the Na^+^/K^+^ pump influence Cl^-^ movement via these routes (Hertz et al., 2015). Cl^-^ ion fluxes have been implicated in astrocyte volume regulation in response to K^+^ in the hippocampus (Larsen and MacAulay, 2017). Therefore, we set out to test whether external Cl^-^ was involved in this phenomenon. Reducing external Cl^-^ from 133.5mM to 7.5mM by substituting NaCl with sodium gluconate, decreased the magnitude of both the decrease in Ca^2+^ (0.9 ± 0.01, p=0.003, n=8, Figure 5A) and the soma swelling caused by a +2.5mM K^+^ challenge (p=0.02, Figure 5B). We then explored the involvement of anion channels and transporters that move Cl^-^. Our first attempt with the broad-spectrum Cl^-^ channel antagonist DIDS (500 μM) was untenable due to volume dysregulation (blebbing) and Ca^2+^ escalation in the brain slice from applying this compound alone (data not shown). However, a different broad-spectrum anion channel antagonist NPPB (100 μM), which targets Volume Regulated Anion Channels (Inoue and Okada, 2007), Ca^2+^-dependent Chloride Channels (Huang et al., 2012) and Max Anion channels (Dutta et al., 2008), was tenable within the time frame of the experiment. While NPPB also caused aberrant cell volume changes across the brain slice at approximately 20min into the application (red brackets, Figure 5E,F), we were able to conduct a 5min NPPB pre-incubation with a +2.5mM [K^+^]_o_ challenge. Interestingly, the drop in astrocyte Ca^2+^ still occurred in NPPB (p=0.96, Figure 5C,E,K), yet the cell swelling was blocked up to 900sec (1.05 ± 0.01, p=0.001, n=8, Figure 5D,F,L), after which time brain slice stability degraded by NPPB treatment.

To test Cl-transporters, we focused on proteins that moved both K^+^ and Cl^-^. For example, KCC moves K^+^ and Cl^-^ out of the cell and KCC1 and KCC3 are detected in astrocytes (Cahoy et al., 2008; Ringel and Plesnila, 2008), whereas NKCC is not (Cahoy et al., 2008). We employed bumetanide, a common NKCC and KCC antagonist that is more potent for NKCC (Payne et al. 2003). Intriguingly, we found that a dose of bumetanide that antagonized both transporters (100 μM) blocked the decrease in astrocyte free Ca^2+^ in response to high [K^+^]_o_ (0.93 ± 0.02, p=0.001, n=8, Figure 5G,I,K), whereas a lower dose that putatively blocks only NKCC (2 μM) had no effect (p=0.85, Figure 5K). Interestingly, high dose bumetanide had little impact on cell swelling caused by high [K^+^]_o_ (p=0.85, Figure 6H,J,L). We also found that bumetanide application itself caused little change to resting astrocyte Ca^2+^ (0.96 ± 0.16, n=7), suggesting that the block of the drop in astrocyte Ca^2+^ by bumetanide was not due to occlusion or a floor effect. With a potential role of KCC transporters, we tested if neural-specific KCC2 could be indirectly involved in this phenomenon that was observed in astrocytes. However, the selective KCC2 blocker VU-0240551 (10 μM) failed to block the drop in astrocyte Ca^2+^ caused by a +2.5mM [K^+^]_o_ challenge (0.82 ± 0.03, p=0.87, n=4, Figure 5K), suggesting that neural KCC2 was not involved.

**Figure 6:**
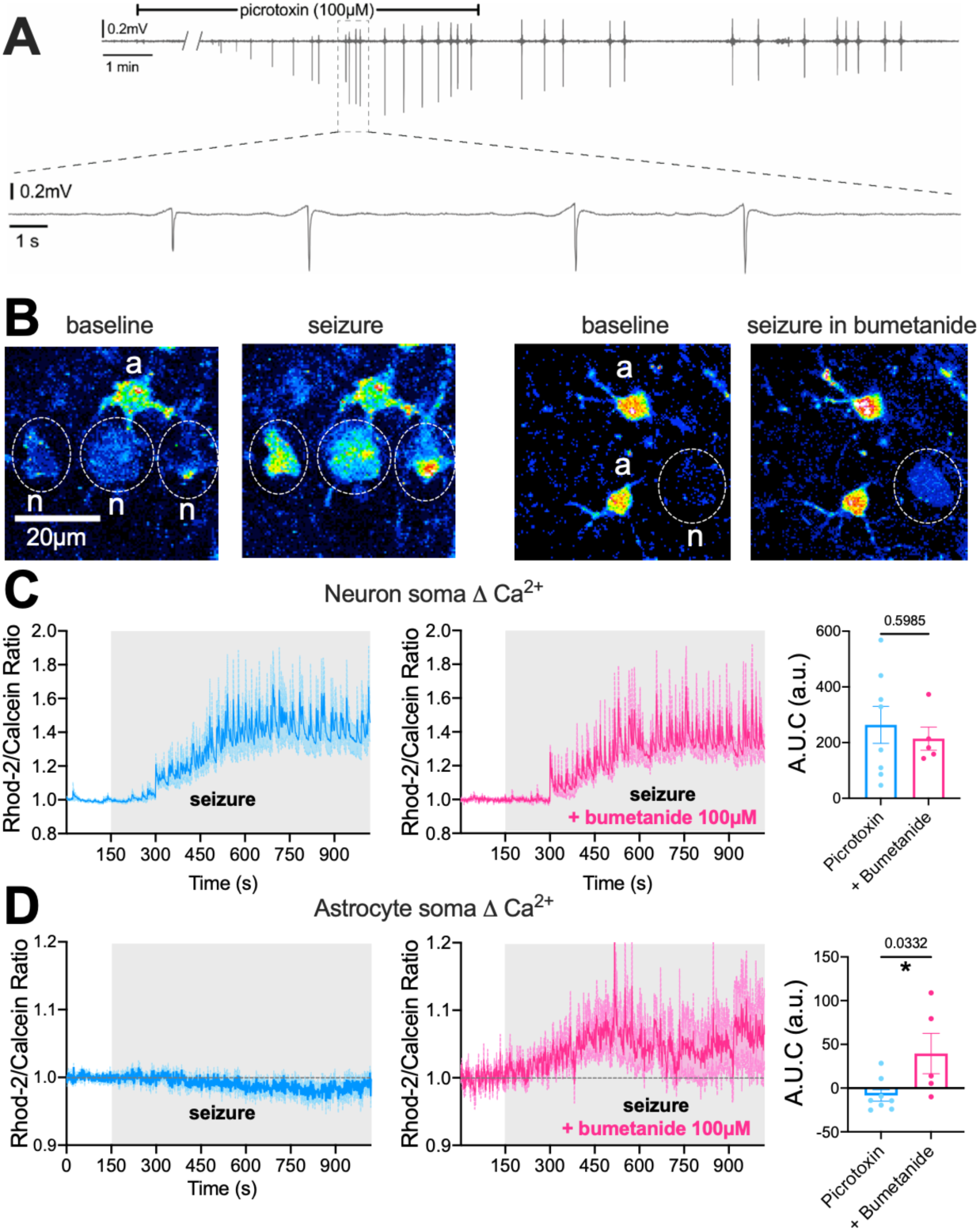
Picrotoxin-induced seizure triggers a bumetanide-sensitive Ca^2+^ drop in cortical astrocytes. **A)** Representative local field potential trace in response to 100μM picrotoxin treatment for 20 min shows repetitive spiking activity. **B**) Pseudo coloured two-photon images of Rhod-2-loaded astrocytes (a) and neurons (n) evoke large increases in neuronal Ca^2+^ and no change in free astrocyte Ca^2+^ to seizure (left), but astrocyte free Ca^2+^ increases in the presence of bumetanide (100 μM). **C-D**) Summary time series of *(C)* neuron soma and *(D)* astrocyte soma Rhod-2/Calcein ratio Ca^2+^ data (average of 2-5 cells/slice) during seizure alone (blue), with bumetanide treatment (magenta) and the comparison of net area under the curve (A.U.C.) of the Ca^2+^ traces. Data is mean +/-SEM, unpaired two-tailed t-tests, * p<0.05.

Collectively, the differential block of NPPB and bumetanide on swelling and Ca^2+^ respectively, strongly suggest each phenomena occurs through distinct mechanisms: K^+^-induced swelling depended on Cl^-^ ion channels and the K^+^-induced Ca^2+^ change depended on potassium and chloride co-transport activity, whereas both phenomena depended on external bicarbonate and external Cl^-^.

### Ictal activity is accompanied with a bumetanide-sensitive reduction of astrocyte Ca^2+^

The synchronous bursting of neuron populations is well known to elevate [K^+^]_o_ (Dreier and Heinemann, 1991; Heinemann et al., 1977). We hypothesised that cortical seizure increases [K^+^]_o_ to a degree that affects astrocyte free [Ca^2+^]_i_ via the same mechanism as seen in our experimental elevation of [K^+^]_o_ by +2.5mM. We chose the GABAa receptor inhibitor picrotoxin for seizure induction as electrical stimulation and depolarizing agents (4-aminopyridine, penicillin, Mg^2+^ free solution) can directly trigger large Ca^2+^ transients in astrocytes, but not GABA_A_-receptor antagonists (Tian et al., 2005). Picrotoxin (100μM) administration for 20 min elicited synchronous neuronal discharges, represented as spikes on the local field potential recording (Fig. 6A) and as simultaneous Ca^2+^ transients with 2-photon imaging in layer 2-3 neuron somata along with a gradual escalation of resting neuronal Ca^2+^ levels (Fig. 6B,C).

Picrotoxin did not change free astrocyte Ca^2+^ before synchronous neuronal discharge occurred (Fig. 6A) and produced only a slight reduction in response to seizure activity (0.9645 ± 0.018)(Fig. 6B,D). Ictal neuronal activity in the presence of the K^+^ Cl^-^ cotransporter blocker bumetanide, however, increased the integrated free Ca^2+^ response curve (area under the curve (A.U.C): 39.55 ± 23 a.u. N=5 slices, 22 cell) compared to the control seizure condition (AUC: - 8.38 ± 6.6 a.u., p=0.0332). Bumetanide is recognized for its antiepileptic properties by modulating GABAergic inhibition (Löscher et al., 2013), so we compared the size of picrotoxin-induced neuronal Ca^2+^ response with or without bumetanide treatment. The integrated Ca^2+^ response of neurons showed no difference with bumetanide treatment (A.U.C: 214 ± 41 a.u. N=5 slices, 22 cells) relative to the control group (A.U.C: 263 ± 66 a.u. N=5 slices, 22 cells, p=0.5985). This indicates that the relatively steady-state free astrocyte Ca^2+^ level is a consequence of a balance between a K-Cl cotransport-mediated decrease and, perhaps, an extracellular glutamate-induced increase of free cytosolic Ca^2+^.

### High [K^+^]_o_ dilates penetrating arterioles in part through an astrocytic mechanism

Elevations in [K^+^]_o_ dilate cerebral arterioles, though most experiments test with ≥8mM. The opening of vascular Kir 2.1 channels is facilitated by high [K^+^]_o_, which leads to hyperpolarization of the membrane potential and vessel relaxation (Longden et al., 2017)(Schubert et al., 2004). High [K^+^]_o_ in the parenchyma may access the perivascular space to initiate vasodilation either directly through limited gaps between adjacent endfeet (Mathiisen et al., 2010) or from K^+^ efflux from endfeet onto the vessel after astrocytes take up excess external K^+^ from around synapses. Elevations in astrocyte Ca^2+^ from strong synaptic activity is thought to initiate K^+^ efflux from endfeet occurs through BK channels (Filosa et al., 2006; Girouard et al., 2010) but not through endfoot K_ir_ channels (Metea et al., 2007). However, a role for K-Cl cotransport in [K^+^]_o_ and [Ca^2+^]_i_ dependent dilation in cerebral penetrating arterioles is unclear. These cotransporters could represent an alternative route for K^+^ efflux from endfeet onto the vessel, or they could initiate vasodilation via changes to the free Ca^2+^ concentration in endfeet, altering the release of vasoactive substances. We found that a +2.5mM [K^+^]_o_ increase dilated, pre-constricted (U-46619 100nM) cerebral penetrating arterioles in brain slices (1.07 ± 0.01, p<0.001, n=13, Figure 7A,B,C), which returned to baseline after high [K^+^]_o_ washout. Notably, bumetanide (100μM) completely blocked the high [K^+^]_o_ induced dilation, resulting in a small vasoconstriction (0.92 ± 0.013, p=0.003, n=8, Figure 7B,C), consistent with an important role for K-Cl cotransport.

**Figure 7:**
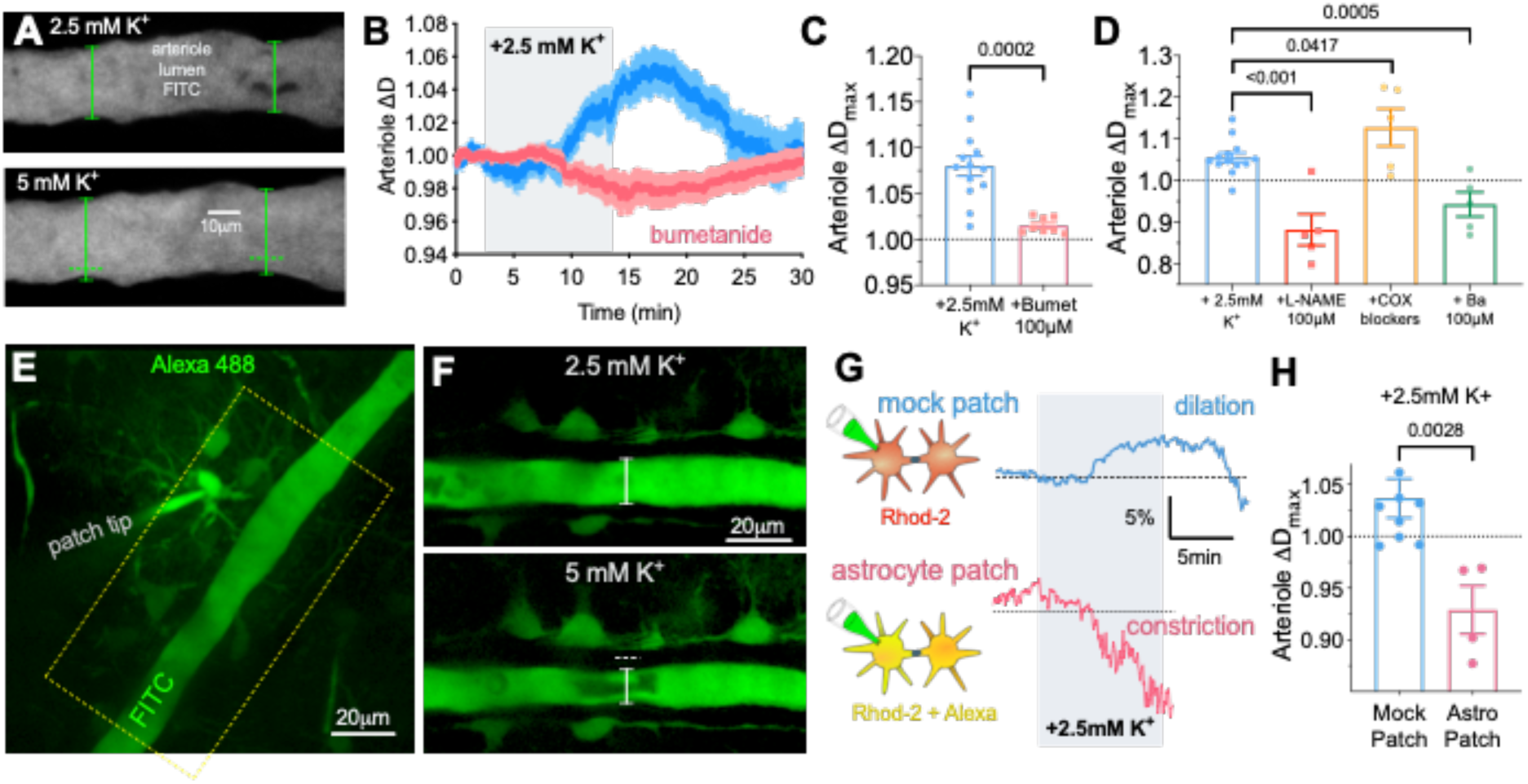
K^+^-mediated vasodilation requires K-Cl co-transport, NO, Kir, and is blocked by astrocyte patching. **A**) Images of the arteriole lumen filled with FITC-dextran showing a vasodilation in response to a +2.5mM increase in external K^+^ in acute slices. **B**) Summary time series data of arteriole diameter in response to +2.5mM K^+^ in the control condition (blue), and in the presence of bumetanide (pink). Bumetanide blocks the high K^+^ mediated vasodilation which instead manifests as a small vasoconstriction. **C**) Summary data of peak arteriole diameter changes related to *B)*. **D**) Arteriole diameter changes at the peak of the summary curve during +2.5mM K+ in the presence of L-NAME, COX-1 (SC560) and COX-2 (SC58125) blockers and Barium. **E**) Two-photon z-stack image showing a patched astrocyte, the coupled astrocytes and perivascular endfeet (Alexa 488) opposed to a penetrating arteriole (FITC-dextran in lumen). **F**) Close up of the arteriole and opposed endfeet loaded with Alexa 488 via the patched astrocyte (not shown, above image plane). A +2.5mM K^+^ increase caused a vasoconstriction when the astrocyte was patched. **G**) Upper: trace of the vasodilation observed in response to high K^+^ in the presence of a ‘mock astrocyte patch’ experiment (pipette sealed/abutted to the cell membrane but no whole-cell). Lower: trace of the vasoconstriction observed in response to high K^+^ in the presence astrocyte network filling with a standard, HEPES buffered internal solution. **H**) Summary data of peak arteriole diameter changes. Data is mean +/-SEM, unpaired two-tailed t-tests, ** p<0.01, *** p<0.001

Testing other classic vasoactive pathways, as expected we found that vasodilation to moderate [K^+^]_o_ elevation was blocked by the K_ir_ blocker Ba^2+^ (100μM) (Figure 7D). The NOS blocker L-NAME (100μM) also prevented the dilation to +2.5mM K^+^ (Figure 7D). However, the COX-1 blocker SC560 (1μM) and the COX-2 blocker SC-58125 (1μM) combined had an opposite effect, potentiating the dilation response to elevated [K^+^]_o_ (Figure 7D), which was likely due to increased arteriole tone from the COX-1 antagonist (Haidey et al., 2021; Rosenegger et al., 2015), increasing the dilatory range.

The block of vasodilation by bumetanide may not necessarily localize to astrocytes, as studies on arterioles outside the brain have described a role for K-Cl cotransport in contractile function (Garneau et al., 2016; Löscher et al., 2013). To test the idea that the influence of high [K^+^]_o_ on the arteriole was conducted, at least in part, through astrocytes, we tested the hypothesis that disrupting normal ion homeostasis in the network of astrocytes surrounding a penetrating arteriole compromises K^+^-induced vasodilation. A common intracellular whole-cell patch solution involving EGTA, low [Cl^-^]_i_ and a HEPES pH buffer, perturbs the natural level and/or movements of Ca^2+^ (Zheng et al., 2015), Cl^-^ (Kyrozis and Reichling, 1995) and HCO_3-_ (Staley and Proctor, 1999). We patched peri-arteriole astrocytes and loaded them with a standard intracellular solution plus Alexa 488 sodium hydrazide (200μM) to visualize the extent of astrocyte network loading and to ensure the solution infiltrated endfeet apposed to the vessel wall. Previously we reported that this intracellular solution had little impact on resting arteriole diameter (Rosenegger et al., 2015). After allowing at least 10min for the astrocyte patch solution to diffuse and equilibrate, we bath applied a +2.5mM [K^+^]_o_ challenge and found that high K^+^-induced dilation was converted into a vasoconstriction (0.93 ± 0.02, n=4, Figure 7D-G). To control for the possibility that the patching process itself caused the switch from vasodilation to vasoconstriction, as opposed to a disruption of the intracellular milieu of astrocytes, we conduced mock patching experiments. Here, rather than achieving astrocyte whole-cell, an on-cell configuration was maintained. Under these conditions, the K^+^-induced vasodilation still occurred (1.04 ± 0.02, n=9, Figure 7F,G), which was different from experiments with whole-cell patch (p=0.007). These data show that antagonizing K-Cl cotransport activity or a general disruption of the internal astrocytic environment, blocks vasodilation to a modest increase in external K^+^.

## Discussion

Here we show that a modest increase or decrease in external [K^+^] bi-directionally and reciprocally controls the resting free Ca^2+^ concentration and cell volume in astrocytes. Resting Ca^2+^ is a recent consideration, given observations that astrocytes appear to have a higher resting Ca^2+^ concentration than neurons *in vitro* and *in vivo* (Shigetomi et al., 2012; 2013b; Zheng et al., 2015) and at least two populations of astrocytes can be designated by different resting Ca^2+^ levels (Zheng et al., 2015). Furthermore, astrocyte resting Ca^2+^ is thought to contribute to 1) constitutive D-serine release (Shigetomi et al., 2013b), 2) basal arteriole tone regulation (Rosenegger et al., 2015; Mehina et al., 2017) and 3) basal release probability at excitatory synapses (Panatier et al., 2011). Evidence shows that the steady-state Ca^2+^ concentration in the cytosol is at least partly dependent on Ca^2+^ influx from outside the cell. This could occur via spontaneous microdomain Ca^2+^ transients, or via constant influx, each through specific ion channels. Some propose a Ca^2+^ influx pathway through TRPA1 channels, regulating both microdomain transients and the baseline Ca^2+^ level (Shigetomi et al., 2012; Shigetomi et al., 2013b). However, others have shown that TRPA1 is not responsible for the microdomain Ca^2+^ transients (Rungta et al., 2016). Though the resting Ca^2+^ level was not directly examined in this latter study, bath applying a Ca^2+^ free external solution plus EGTA decreased the baseline Ca^2+^ level in flou-4 patch loaded astrocytes (Rungta et al., 2016). Our data is consistent with the idea that a modest elevation in [K^+^]_o_ decreased ongoing Ca^2+^ influx across the plasma membrane, but not via decreased release from internal Ca^2+^ stores. We expect that the decrease in cytosolic free Ca^2+^ occurred in Ca^2+^ free solution, as shown by others previously, which could not be decreased further when [K^+^]_o_ became elevated. It remains unclear how free cytosolic Ca^2+^ decreases due to decreased influx. The bidirectional nature of the Ca^2+^ change, and the well-documented influence of external K^+^ on controlling astrocyte resting membrane potential, leads us to speculate that a change in membrane voltage is an aspect to the phenomenon. Small increases in external K^+^ of 1mM will shift E_K_ ∼9mV, which, after a subsequent depolarizing shift in membrane potential, would lessen the driving force for the inward movement of Ca^2+^. From this, one might expect that we could block the drop in Ca^2+^ by clamping membrane potential at a hyperpolarized value, such as in our FLIM experiments.

However, astrocytes are poorly voltage clamped given their high resting permeability to K^+^ and very low input resistance (Steinhauser et al., 1992; Ma et al., 2014). What type of K-Cl co-transport is involved in the K+-mediated changes to astrocyte free Ca^2+^? Astrocyte transcriptome evidence from the neocortex shows no detectable expression of NKCCs in astrocytes, whereas KCC1 and KCC3 are clearly transcribed (Cahoy et al., 2008; Zhang et al., 2014). Importantly, though bumetanide is typically a NKCC blocker, it does block KCC1 at higher doses (Gillen and Forbush, 1999). We found no block of the effect with 2 μM bumetanide, which should only block NKCCs, whereas 100 μM was highly effective. That low Na^+^ external solution did not block the drop in Ca^2+^ to high K^+^ also argues against the involvement of NKCCs. These observations lead us to support a role for KCCs over NKCCs in our described effects. However, two limitations are: 1) we were only able to decrease [Na^+^]_o_ to 26.25 mM and the difference in the Ca^2+^ drop to control was nearly significant (p=0.06). A further decrease in external Na^+^ may have revealed a significant reduction in the Ca^2+^ drop. 2) As 2 μM bumetanide is ∼twice the IC50 for NKCC, a higher dose may be necessary in acute slices to achieve a block. Furthermore, 100μM is expected to block KCCs, yet, this would be on the lower end of the dose response curve. Thus, further experiments knocking down KCCs vs NKCCs in astrocytes is warranted.

Though K_ir_ 4.1 on astrocytes contributes to K^+^ buffering, and the K^+^ influx could increase KCC activity, we found no effect of 100μM Ba^2+^ on the K^+^-induced decrease in astrocyte Ca^2+^. Previous work observed that decreasing external K^+^ can increase Ca^2+^ via a BaCl_2_ sensitive pathway in cultured astrocytes and *in situ* (Dallwig et al., 2000). While this is analogous to the effects we observed with a 1mM decrease in [K^+^]_o_, this work employed a larger unphysiological drop than we explored: moving from 5mM to 0.2 or 0.4 mM [K^+^]_o_. If not inward rectifiers, what invokes K-Cl co-transport? High [K^+^]_o_ will increase Na^+^/K^+^ ATPase activity, increasing intracellular [K^+^], which could subsequently increase the efflux of K^+^ and Cl^-^ via the co-transporters. However, blocking this crucial ATPase to test its role in our effect is untenable in brain slices at the micron-level due to robust cell volume changes to the antagonist ouabain (Joshi and Andrew, 2001). Alternatively, high external K^+^ would also limit KCC co-transport activity by acting against the outward movement of K+ and Clions. The cumulative effect on K+ and Cl-transport activity would depend on the relative expression, the kinetics of each transport system, as well as their respective locations in the cell. For example, if KCCs were targeted to the vascular interface of endfeet and thus sheltered directly from parenchymal elevations in K^+^. Undoubtedly, changing K-Cl cotransport activity will affect the intracellular K+ and Cl-concentrations, and potentially membrane potential due to effects on other transport mechanisms or ion channels. For instance, bumetanide itself influences membrane potential in kidney cells (Wang et al., 2013) and skeletal muscle (van Mil et al., 1997) and bumetanide can prevent membrane potential changes to osmotic stimuli (van Mil et al., 1997).

We demonstrated that during cortical seizure, K^+^ buffering by astrocytes drives a K-Cl cotransport-mediated reduction of free Ca^2+^. This reduction, however, is counteracted by other factors (likely glutamate) rising free Ca^2+^ level to produce an overall steady Ca^2+^ concentration in astrocytes. The elevation of astrocyte free Ca^2+^ during seizures was shown to promote and maintain ictal activity (Gómez-Gonzalo et al., 2010). The K-Cl-cotransport mediated reduction of astrocytic Ca^2+^ likely exerts an anti-ictal effect when seizure elevates [K^+^]_o_. This phenomenon needs further exploration to confirm the anti-epileptic property of K^+^-induced astrocyte Ca^2+^ drop in other seizure models and *in vivo*.

It is important to consider whether the decrease in astrocyte Ca^2+^ caused by a moderate elevation in [K^+^]_o_ was itself sufficient to cause high K^+^-induced vasodilation. From one perspective, our data argue against the sufficiency of astrocyte Ca^2+^ because although patching a peri-arteriole astrocyte blocks K^+^-induced vasodilation, Ca^2+^ is likely still decreasing in astrocytes. This was clearly the case given our FLIM Ca^2+^ measures, which introduced OGB-1 into astrocytes via the patch pipette. In this condition, the Ca^2+^ drop could be detected in all compartments of the patched cell, including peri-vascular endfeet. Thus, the patch could have disrupted a different ion such as Cl^-^ or HCO_3-_ to change the response of the vessel. From another perspective, the resting Ca^2+^ level is significantly higher in the patched cell compared to coupled cells. Thus, even though Ca^2+^ still decreased while patched in response to high [K^+^]_o_, the absolute level of Ca^2+^ reached was not as low as un-patched astrocytes (∼75nM compared to ∼40nM). This could affect the type or amount of vaso-active messenger being released from astrocytes, which could explain why we observed a vasoconstriction to high K^+^ instead of a vasodilation while the peri-vascular astrocyte was patched. It is important to note that we previously observed a vasoconstriction when astrocyte Ca^2+^ was lowered to <25nM via a BAPTA loaded patch pipette (Rosenegger et al., 2015; Haidey et al., 2021) and long lasting increases in arteriole tole (vasoconstriction) to glutamate receptor activation that were associated with a decrease in endfoot Ca^2+^. These are opposite to what we observed with moderate high [K^+^]_o_, in which a free Ca^2+^ decrease was associated with a vasodilation. Key differences could be 1) the absolute level of Ca^2+^ achieved with each manipulation and 2) the microdomains where the effects were occurring. For example, BAPTA affects the entire cytosol indiscriminately, whereas K^+^ and glutamate will have microdomain effects for where transporters, receptors and vasoactive enzymes are located. Measuring such changes in free Ca^2+^ with a freely diffusible, cell-wide Ca^2+^ indicator cannot distinguish between different mechanisms occurring in unique microdomains.

## METHODS

### EXPERIMENTAL MODEL AND SUBJECT DETAILS

All animal procedures were approved by the Animal Care and Use Committee of the University of Calgary (protocols AC15-0053 and AC15-0133). All studies were performed on either male Sprague Dawley rats between P23 to P30, or male C57BL/6 mice between P30 to P60 (Charles River, Wilmington, MA, USA). Animals were kept on a standard 12 hour dark 12 hour light cycle and had *ad libitum* access to food and water. Animals were group housed until head-bar installation for in vivo experiments. The experimenter was not blinded to any treatment.

## METHOD DETAILS

### Acute Brain Slice Preparation

Care and use of animals for this project was carried out in accordance with approved guidelines set forth by the University of Calgary Animal Care and Use Committee. Male Sprague Dawley rats (P23-P30) were deeply anaesthetized with gaseous isofluorane (5%) and then decapitated using a rodent guillotine. The brain was rapidly and surgically removed, then submerged for ∼2 minutes in ice-cold slicing solution containing (in mM): 119.9 N-methyl-D-glucamine, 2.5 KCl, 25 NaHCO3, 1.0 CaCl2-2H2O, 6.9 MgCl2-6H2O, 1.4 NaH2PO4-H2O, and 20 D-glucose. The brain was then Krazy Glued onto a vibratome tray (Leica Instruments, VT1200S) and then resubmerged in ice-cold slicing solution. Acute coronal slices were prepared from the somatosensory cortex (400 μm thick). The slices were incubated for 45 minutes at 33ºC in a recovery chamber filled with artificial cerebrospinal fluid (ACSF) containing (in mM): 126 NaCl, 2.5 KCl, 25 NaHCO3, 1.3 CaCl2-2H2O, 1.2 MgCl2-6H2O, 1.25 NaH2PO4-H2O, 10 glucose. At all steps of tissue preparation and for all experiments using bicarbonate buffered ACSF, the brain/slices were continuously bubbled with carbogen (95% oxygen, 5% carbon dioxide).

### Contents of Various ACSF Solutions

High K+ ACSF was bath applied in order to raise external [K+] to 5.0 mM. In order to keep high K+ ACSF solutions iso-osmotic with regular ACSF, the solutions were prepared by adding additional KCl to achieve 5.0 mM while an equimolar amount of NaCl was removed. This solution contained (in mM): 123 NaCl, 5 KCl, 25 NaHCO3, 1.3 CaCl2-2H2O, 1.2 MgCl2-6H2O, 1.25 NaH2PO4-H2O, 10 glucose. For all experiments using bicarbonate buffered ACSF, the brain slices were continuously bubbled with carbogen (95% oxygen, 5% carbon dioxide).

In bicarbonate-free experiments, slices will be transferred into 4-(2-hydroyethyl)-1-piperazineethanesulfornic acid (HEPES) buffered ACSF after the 45 minute incubation (mentioned above) in bicarbonate buffered ACSF. The HEPES buffered ACSF contains (in mM): 142 NaCl, 2.5 KCl, 10 HEPES, 1.3 CaCl2-2H2O, 1.2 MgCl2-6H2O, 1.25 NaH2PO4-H2O, 10 glucose. The HEPES buffered ACSF used to increase extracellular [K+] to 5.0 mM contains (in mM): 139 NaCl, 5 KCl, 10 HEPES, 1.3 CaCl2-2H2O, 1.2 MgCl2-6H2O, 1.25 NaH2PO4-H2O, 10 glucose. For all experiments using HEPES buffered ACSF, the brain slices were continuously bubbled with 100% oxygen and pH corrected with NaOH to 7.4.

For experiments substituting Na+, NaCl was replaced with choline chloride (C5H14ClNO) in order to maintain Cl-levels and osmolarity. This solution contained (in mM): 126 C5H14ClNO, 2.5 KCl, 25 NaHCO3, 1.3 CaCl2-2H2O, 1.2 MgCl2-6H2O, 1.25 NaH2PO4-H2O, 10 glucose. The ACSF used to increase extracellular [K+] to 5.0 mM with this substitution contained (in mM): 123 C5H14ClNO, 5 KCl, 25 NaHCO3, 1.3 CaCl2-2H2O, 1.2 MgCl2-6H2O, 1.25 NaH2PO4-H2O, 10 glucose. Brain slices were continuously bubbled with carbogen (95% oxygen, 5% carbon dioxide).

For experiments substituting Cl-, NaCl was replaced with sodium gluconate (C6H11NaO7) in order to maintain Na+ levels and osmolarity. This solution contained (in mM): 126 C6H11NaO7, 2.5 KCl, 25 NaHCO3, 1.3 CaCl2-2H2O, 1.2 MgCl2-6H2O, 1.25 NaH2PO4-H2O, 10 glucose. The ACSF used to increase extracellular [K+] to 5.0 mM with this substitution contained (in mM): 123 C6H11NaO7, 5 KCl, 25 NaHCO3, 1.3 CaCl2-2H2O, 1.2 MgCl2-6H2O, 1.25 NaH2PO4-H2O, 10 glucose. Brain slices were continuously bubbled with carbogen (95% oxygen, 5% carbon dioxide).

In experiments removing external Ca2+, Ca2+ free ACSF external solutions were made. This solution contained (in mM): 126 NaCl, 2.5 KCl, 25 NaHCO3, 1.2 MgCl2-6H2O, 1.25 NaH2PO4-H2O, 10 glucose. The Ca2+-free solution used to increase extracellular [K+] to 5.0 mM contained (in mM): 123 NaCl, 5 KCl, 25 NaHCO3, 1.2 MgCl2-6H2O, 1.25 NaH2PO4-H2O, 10 glucose. Brain slices were continuously bubbled with carbogen (95% oxygen, 5% carbon dioxide).

### Loading Dyes in Astrocytes, Neurons, and Arterioles

After recovery from slicing, the tissue was incubated in a 3 mL well at 33ºC for 45 minutes with the calcium indicator Rhodamine-2 acetoxymethyl ester (Rhod-2 AM) (15 μM) and the morphological dye Calcein Green AM (17 μM) (Biotium, Fremont, CA, USA); 0.2% DMSO; 0.006% Pluronic Acid; 0.0002% Cremophore EL. During incubation in the small well, slices received very fine carbogen bubbling using a flexible 34 gauge pipette tip (WPI, Sarasota, FL, USA). This dual labelling allowed us to take a ratio of the changes in the fluorescence of the calcium indicator over that of the changes in the morphological dye. Astrocytes were identified by their bright uptake of Rhod-2 AM and by their perivascular endfeet (Simard et al., 2003; Mulligan and MacVicar, 2004).

For experiments that examined parenchymal arterioles, the animal received a tail vein injection of 15 mg Fluorescein Isothiocyanate (FITC) dextran (2000 kDa, Sigma Aldrich) dissolved in 300 μL of lactated Ringer’s solution under isoflurane anesthesia (5% induction, 2% maintenance) before decapitation. Luminal FITC-dextran permitted visualization of the brain microvasculature and we quantified arteriole diameter as a change in lumen area. For an experiment, a given slice was transferred to a superfusion chamber on the rig and was perfused using a pressure driven (carbogen) solution delivery system at ∼1.0 mL per minute, maintained at room temperature. For experiments examining arterioles, the thromboxane A2 analog U46619 (100 nM) was present continually in the bath to provide constant artificial tone, as was done previously (Institoris et al., 2015).

### Two-Photon Fluorescence Microscopy

Slices were imaged using a custom built two-photon microscope (Rosenegger et al., 2014) fed by a Ti:Sapph laser source (Coherent Ultra II, ∼4 W average output at 800 nm, ∼80 MHz). Image data were acquired using MatLab (2013) running an open source scanning microscope control software called ScanImage (version 3.81, HHMI/Janelia Farms) (Pologruto et al., 2003). Imaging was performed at an excitation wavelength of 850 nm for Rhod-2/Calcein experiments. The microscope was equipped with a primary dichroic mirror at 695 nm and green and red fluorescence was split and filtered using a secondary dichroic at 560 nm and two bandpass emission filters: 525-40 nm and 605-70 nm (Chroma Technologies). Time series images were acquired at 0.98 Hz with a pixel density of 512 by 512 and a field of view size of ∼150 μm. Imaging used an Olympus 40x water dipping objective lens (NA 0.8, WD 3.0 mm).

### Fluorescence Lifetime Imaging Microscopy (FLIM)

Time resolved fluorescent lifetime imaging: Acute slices from Sprague Dawley rats (P21-28) were incubated in 1 μM SR101 to load astrocytes for identification for whole-cell patch clamp. The pipette recording solution consisted of (in mM): 113 K-Gluconate, 3 KCl, 8 Na-Gluconate, 2 MgCl2, 4 K2ATP, 0.3 NaGTP, 10 HEPES, 1 EGTA, 0.23 CaCl2, 0.2 OGB-1 Hexapotassium salt (Thermo Fischer), pH and osmolarity adjusted to 7.25 (with KOH) and 290 mOsm, respectively.

SR101 positive cells were whole-cell patch clamped and were dialyzed for at least 20 min prior to imaging to permit sufficient OGB-1 dye dialysis to gap-junctionally coupled neighbours. FLIM images were acquired with a Zeiss LSM 7 imaging system retrofitted with FLIM hardware module (SPC-150) from Becker Hickl. OGB-1 was imaged at 800nm using a tunable pulsed 2-photon Ti:Sa femtosecond laser (Coherent) with a 80MHz repetition rate and a 20X water immersion objective (NA=1). Individual images were acquired at 256×256 (x,y) pixels, 32 frame scans per image, with images acquired in 30s intervals to avoid phototoxicity. Lifetime decay data from individual pixels were binned with neighbouring pixels by a factor of 2 to ensure at least 10 photon counts were present at the end of each trial and ensuring proper exponential fitting. The lifetime data were fit based on a 2-component multiexponential decay calculation using Becker & Hickl’s SPCImage (version 6.5) software based on goodness of fit. Raw data are presented as ‘Tau mean’ (τmean), encompassing the weighted contributions of each component of the full decay curve and is expressed as: τmean =α1τ1 + α2τ2, where α1 and α2 represent the fractional relative intensities (i.e. α1+α2=1) of the respective tau components τ1 and τ2. Mean lifetime was used over ‘intensity-weighted average lifetime’ (i.e. decay components weighted by intensity integrals) as τmean calculations are more sensitive to changes at fast lifetimes and therefore low Ca2+ concentrations (Becker, 2012). Calculated τmean data were compared to their respective Ca2+ concentrations via in vitro calibration (below).

### *In vitro* OGB-1 Lifetime Calibration

OGB-1 calibration was performed in sealed recording electrodes at 33ºC similar to that described previously (Zheng et al., 2015). Electrodes were filled with an electrophysiological recording solution consistent of (in mM): 93 K-Gluconate, 8 Na-Gluconate, 2 MgCl_2_, 4 K_2_ATP, 0.3 NaGTP, 10 HEPES, 10 EGTA, 0.2 OGB-1 hexapotassium salt. 1.17-20,000nM of free Ca^2+^ (added as CaCl_2_) was calculated using Webmaxc Standard (http://web.stanford.edu/~cpatton/webmaxcS.htm) and balanced accordingly with KCl and pH adjusted to pH=7.25 with KOH. Note that OGB-1 lifetimes are not affected by changes in pH, temperature, or viscosity (Zheng et al., 2015). Calibration data were fit to a four-variable logistic (sigmoid) function: Y=-4.87*10^6+(3300+4.87*10^6)/(1+10^((2.075-X)*1.573)), where Y = Tau(mean) and X= calcium concentration.

### Enriched Environment

For the enriched animal experiments, 7 Sprague Dawley rats (P21 to P42-45) lived communally in an enriched environment, which was approximately 2 feet x 1.5 feet x 1.5 feet in size for 3 weeks. Within this environment were: a running wheel, tunnels, ladders, hammocks, shelters, and readily accessible food and water.

### Astrocyte Patch-Clamp

To be selected for patch-clamp, astrocytes were required to be located between ∼25 to 40 μm below the surface of the slice and within ∼30 to 50 μm from the arteriole of interest. A Giga-Ohm seal was maintained for 5 minutes, followed by a whole-cell configuration with 15 minutes of astrocyte filling for adequate diffusion of the internal solution into the astrocyte network.

AlexaFluor-488 sodium hydrazide (100-200 μM) was included in the intracellular solution in order to visualize the extent of solution diffusion throughout the astrocyte network. The intracellular solution also contained (in mM): 108 potassium gluconate, 8 KCl, 8 sodium gluconate, 2 MgCl_2_, 10 HEPES, 0.1 potassium EGTA, 4 ATP, and 0.3 sodium GTP. Moreover, the solution was corrected for osmolarity to ∼285 mosmol and corrected for pH with KOH to 7.2. The astrocyte cell type was confirmed by the following: a low input resistance (10-20 MΩ), extensive dye transfer between coupled cells via gap junctions, and visibly loaded endfeet apposed to microvasculature.

### Local field potential measurement

A 3-5 Mega-Ohm pipette, filled with ACSF, was lowered ∼50μm deep in layer 2-3 of the neocortical slice. Local field potential was detected with a Multiclamp 700B amplifier (Molecular Devices), digitized by and Axon Instruments digitizer (1550) and acquired with Clampex version 10 software (Molecular Devices) in gap-free mode, sampled at 10kHz and lowpass fileted at 1kHz using a Bessel filter. Post hoc a high pass 1-50Hz frequency filter was applied to the trace data to identify bursting activity.

### *In vivo* surgery and two-photon imaging

Male, P40-60 c57bl/6 mice (N=3) were used. First, under isoflurane anesthesia (induction 4%, maintenance 1.5-2%) and pain control (buprenorphen 0.05mg/kg) a light (0.5 g) metal headbar was installed on the occipital bone under aseptic conditions with a three component dental glue (C&B Metabond, Parkell Inc, NY, USA) and dental cement (OrthoJet Acrylic Resin, Land Dental MFG. CO., Inc., IL, USA). A cement wall was mounted around the right parietal bone forming a well around the somatosensory area. A blunt piece of an 18G needle was implanted in the medial side of the well to allow for the superfusion of artificial cerebrospinal fluid (aCSF). When the cement was cured, a 2mm in diameter hole was drilled with a center 3mm lateral and 2mm posterior to the Bregma. A custom-made circular glass coverslip was superglued over the craniotomy. This 3mm-wide coverslip contained 2x 600μm-wide circular holes located off center. Next, the dura under the holes was gently removed under continuous superfusion of a HEPES-based aCSF (in mM: 5 KCl, 142 NaCl, 10 glucose, 10 HEPES, 3.1 CaCl2, 1.3 MgCl_2_, pH 7.4) bubbled with 100% O_2_. Rhod-2 AM (30 μM in aCSF, dissolved in 0.2% DMSO; 0.006% Pluronic Acid; 0.0002% Cremophore EL) was incubated on the brain surface for 45 min. Animals were then injected with a maintenance dose of buprenorphen (0.02mg/kg) and a low dose of subcutaneous chlorprothixene (0.5-1 mg/kg body weight) to transition anesthesia to light sedation as isoflurane was discontinued (Bonder et al., 2014 J Neurosci). Subsequently, 0.2 mL 5% FITC dextran solution was injected into the tail vein. The animals were then transferred to the imaging rig on a heating pad and head-fixed for 2-photon imaging. Temperature was monitored with a rectal thermometer and was set to 36ºC. The cranial window was continuously superfused with a bicarbonate based aCSF (in mM: 2.5 KCl, 126 NaCl, 25 NaHCO_3_, 1.3 CaCl_2_, 1.2 MgCl_2_, 1.25 NaH_2_PO_4_, and 10 glucose) bubbled with carbogen (95% O_2_, 5% CO_2_) at a rate of 2ml/min. Imaging started ∼30 min later when the animals were awake and stationary but responsive to touch or startle. Mice were continuously monitored by an infrared camera and an LED light. Additional chlorprothixene (0.2-0.4 mg/kg) was injected if any movement was detected. A 16x (Nikon, 0.9NA) water-immersion objective was positioned square to the surface of the window. Imaging was performed at 0.5 Hz over one of the open holes 40-80 μm below the surface. Rhod-2 AM only loaded astrocytes in mice *in vivo* (Tran et al., 2018 Neuron). After 5 min of baseline recording, standard 2.5 mM K^+^-containing aCSF was switched to a 10 mM K^+^ -based aCSF (in mM: 10 KCl, 118.5 NaCl, 25 NaHCO_3_, 1.3 CaCl_2_, 1.2 MgCl_2_, 1.25 NaH_2_PO_4_, and 10 glucose) for 20 min, where Na^+^ was replaced by the excess K^+^ to maintain osmolality.

## Acknowledgements

The Canada Institutes of Health Research (CIHR) supported this study (FDN-148471). Canada Research Chairs (CRC) supported GRG. SS, EM, KG and JH were supported by the Hotchkiss Brain Institute. EM was additionally supported by CIHR. WN is supported by CIHR and Banting Fellowships, BAM is supported by CRC and CIHR (FDN148397). We acknowledge the developers and distributers of ScanImage software through the HHMI/Janelia Farms Open Source License.

## QUANTIFICATION AND STATISTICAL ANALYSIS

Rhod-2 and Calcein trace data were independently normalized (F = F1/F0) – where F is fluorescence, 1 is any given time point and 0 is an average baseline value – followed by a ratio of the two (F_Rhod-2_/F_Calcein_). Regions of interest (astrocyte somata, major process or endfeet) were chosen manually and adjusted in xy position throughout the time series manually if required.

Quantification of arteriole tone changes in brain slices were performed in ImageJ. The arteriole lumen was loaded with FITC-dextran and the luminal area was calculated in every frame for a section of arteriole representative of vessel’s tone change.

### Data Collection and Statistics

In a given experiment, if more than one astrocyte was imaged in the field of view (typical) the normalized ratios from each were averaged together for an ‘n’ of 1. Thus, each experiment, conducted on an independent brain slice constituted a statistical ‘n’ and all data sets used at least N=3 animals. We plotted the average decrease (mean +/-SEM) in the Rhod-2/Calcein ratio for the entire time series and also quantified the peak decrease of individual experiments for bar graphs. Imaging data were stored on a computer for off-line analysis using ImageJ and Graphpad Prism (Version 6). Experimental values are presented as mean ± SEM; statistical analyses were performed using two-tailed student’s t-test (paired or unpaired as appropriate) or a one-way ANOVA when comparing multiple groups. Values of p<0.05 were accepted as statistically significant (* = p<0.05, ** = p<0.01, *** = p<0.001, **** = p<0.0001).

## KEY RESOURCES TABLE

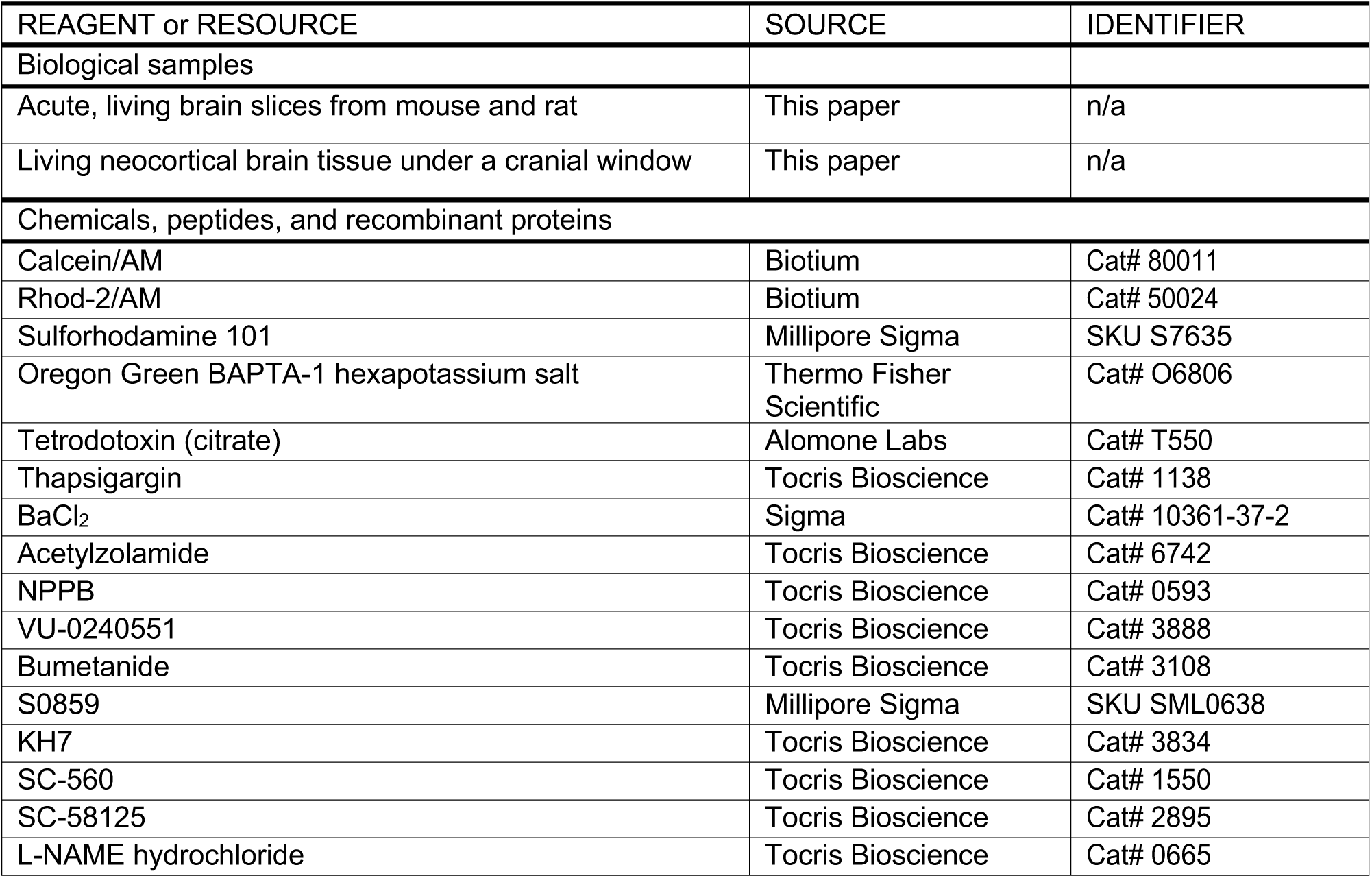
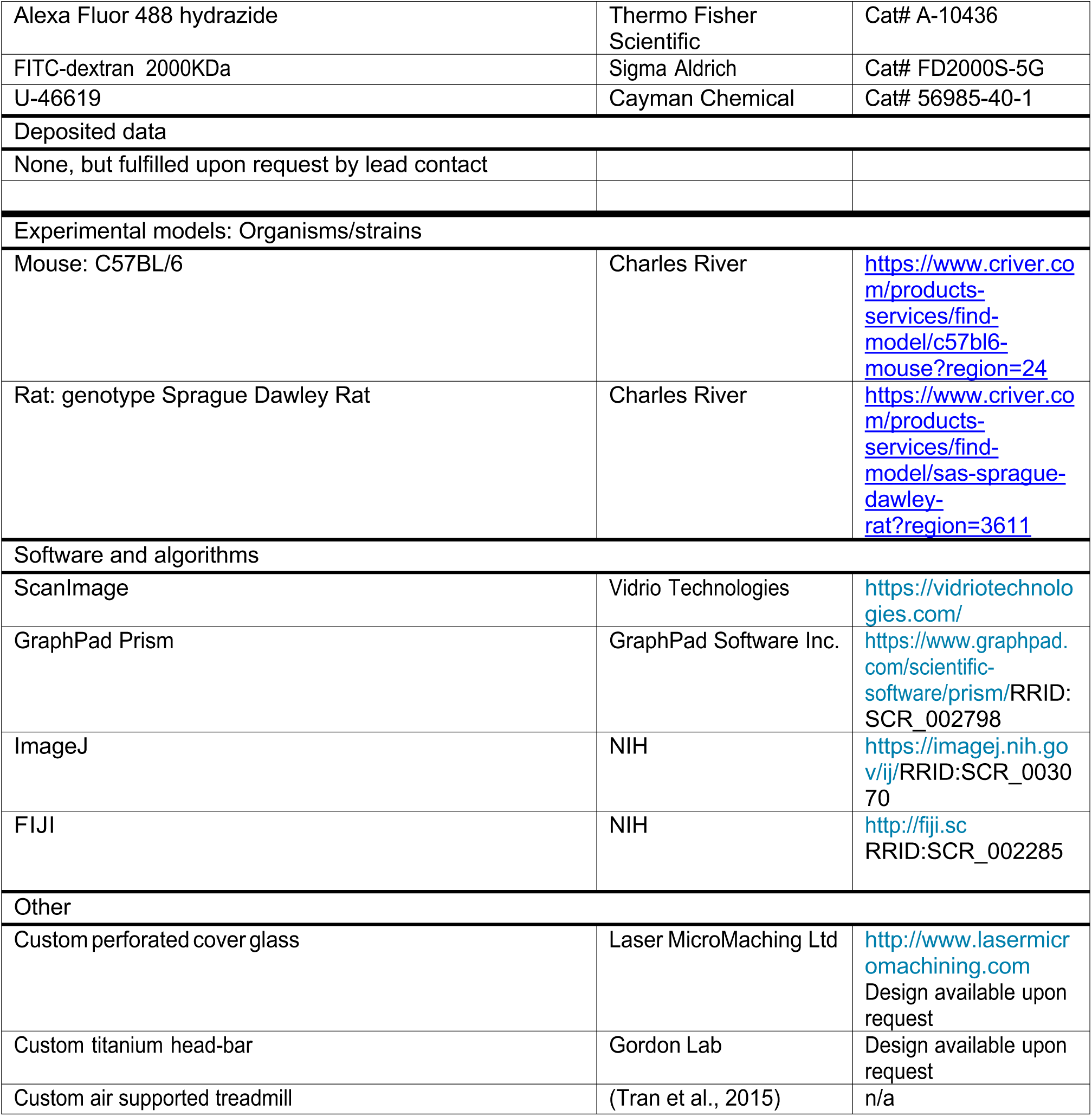

| REAGENT or RESOURCE | SOURCE | IDENTIFIER |
| --- | --- | --- |
| Biological samples |  |  |
| Acute, living brain slices from mouse and rat | This paper | n/a |
| Living neocortical brain tissue under a cranial window | This paper | n/a |
| Chemicals, peptides, and recombinant proteins |  |  |
| Calcein/AM | Biotium | Cat# 80011 |
| Rhod-2/AM | Biotium | Cat# 50024 |
| Sulforhodamine 101 | Millipore Sigma | SKU S7635 |
| Oregon Green BAPTA-1 hexapotassium salt | Thermo Fisher Scientific | Cat# 06806 |
| Tetrodotoxin (citrate) | Alomone Labs | Cat# T550 |
| Thapsigargin | Tocris Bioscience | Cat# 1138 |
| BaCl<sub>2</sub> | Sigma | Cat# 10361-37-2 |
| Acetylzolamide | Tocris Bioscience | Cat# 6742 |
| NPPB | Tocris Bioscience | Cat# 0593 |
| VU-0240551 | Tocris Bioscience | Cat# 3888 |
| Bumetanide | Tocris Bioscience | Cat# 3108 |
| S0859 | Millipore Sigma | SKU SML0638 |
| KH7 | Tocris Bioscience | Cat# 3834 |
| SC-560 | Tocris Bioscience | Cat# 1550 |
| SC-58125 | Tocris Bioscience | Cat# 2895 |
| L-NAME hydrochloride | Tocris Bioscience | Cat# 0665 |
| Alexa Fluor 488 hydrazide | Thermo Fisher Scientific | Cat# A-10436 |
| FITC-dextran 2000KDa | Sigma Aldrich | Cat# FD2000S-5G |
| U-46619 | Cayman Chemical | Cat# 56985-40-1 |
| Deposited data |  |  |
| None, but fulfilled upon request by lead contact |  |  |
| Experimental models: Organisms/strains |  |  |
| Mouse: C57BL/6 | Charles River | https://www.criver.com/products-services/find-model/c57bl6-mouse?region=24 |
| Rat: genotype Sprague Dawley Rat | Charles River | https://www.criver.com/products-services/find-model/sas-sprague-dawley-rat?region=3611 |
| Software and algorithms |  |  |
| ScanImage | Vidrio Technologies | https://vidriotechnologies.com/ |
| GraphPad Prism | GraphPad Software Inc. | https://www.graphpad.com/scientific-software/prism/RRID:SCR_002798 |
| ImageJ | NIH | https://imagej.nih.gov/ij/RRID:SCR_003070 |
| FIJI | NIH | http://fiji.sc<br>RRID:SCR_002285 |
| Other |  |  |
| Custom perforated cover glass | Laser MicroMaching Ltd | http://www.lasermicromachining.com<br>Design available upon request |
| Custom titanium head-bar | Gordon Lab | Design available upon request |
| Custom air supported treadmill | (Tran et al., 2015) | n/a |

## Supplementary text and figures

**Supplementary Figure 1:**
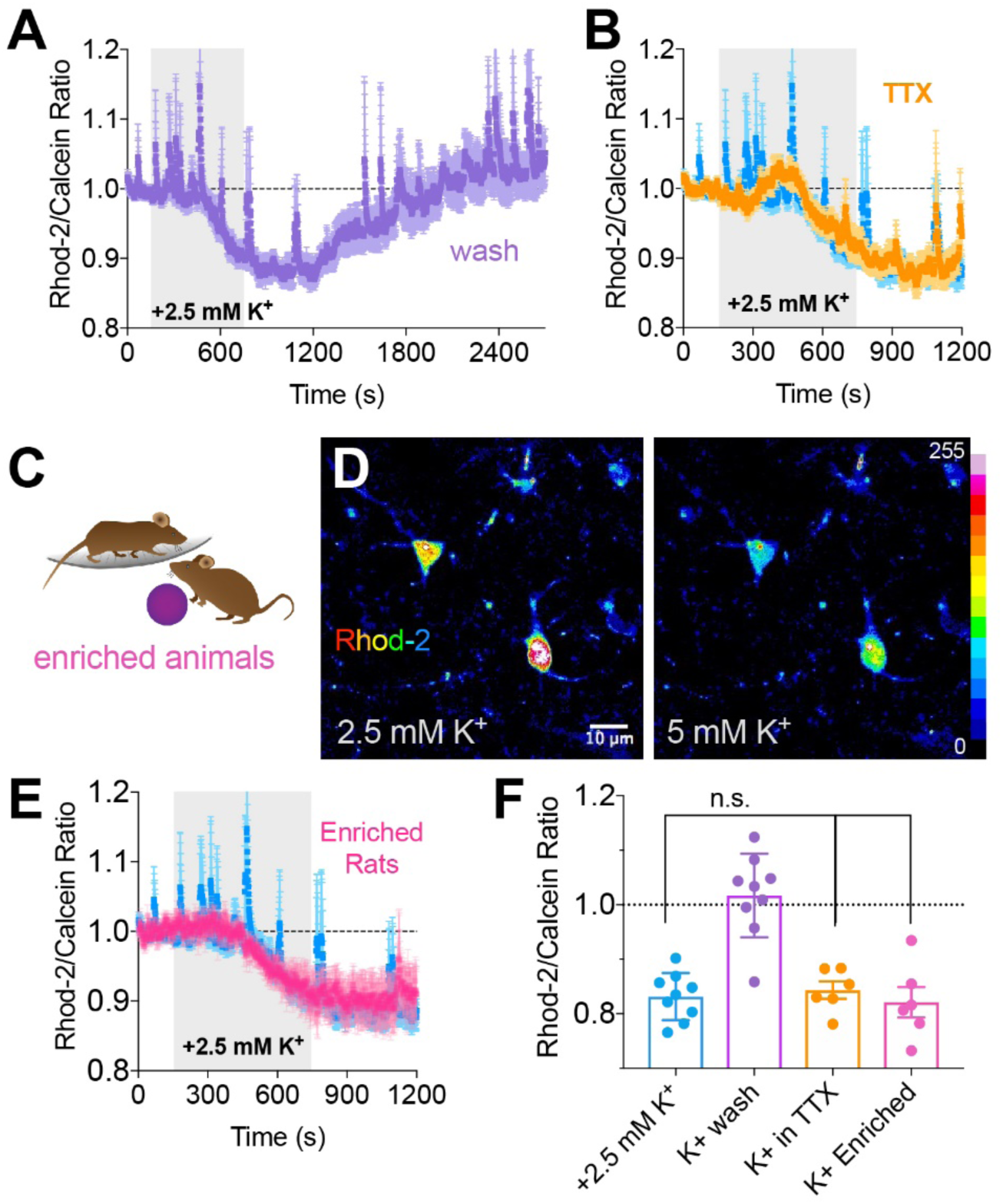
The high K^+^ mediated decrease in astrocyte Ca^2+^ neither exhibits a reliance on neural firing nor displays plastic changes. **A**) Average summary time series Rhod-2/Calcein ratio data showing that the decrease in astrocyte Ca^2+^ caused by a +2.5mM K^+^ challenge recovers completely upon return to the baseline K^+^ level. **B**) The decrease in astrocyte Ca^2+^ caused by high K^+^ still occurs when action potential signaling is blocked by TTX. **C**) Cartoon depicting animals that received 3 weeks of enrichment. **D**) Pseudo coloured images of Rhod-2 astrocytes from enriched animals showing the decrease in Ca^2+^ signal in response to a K^+^ challenge. **E**) Average summary time series of the Rhod-2/Calcein ratio from enriched animals in response to high K^+^ (pink) compared to animals under standard housing (blue). **F**) Summary data of the maximal decrease in astrocyte Ca^2+^ in each experiment. Data is mean +/-SEM, ‘TTX’ and ‘Enriched’ are unpaired two-tailed t-tests to control condition.

**Supplementary Figure 2:**
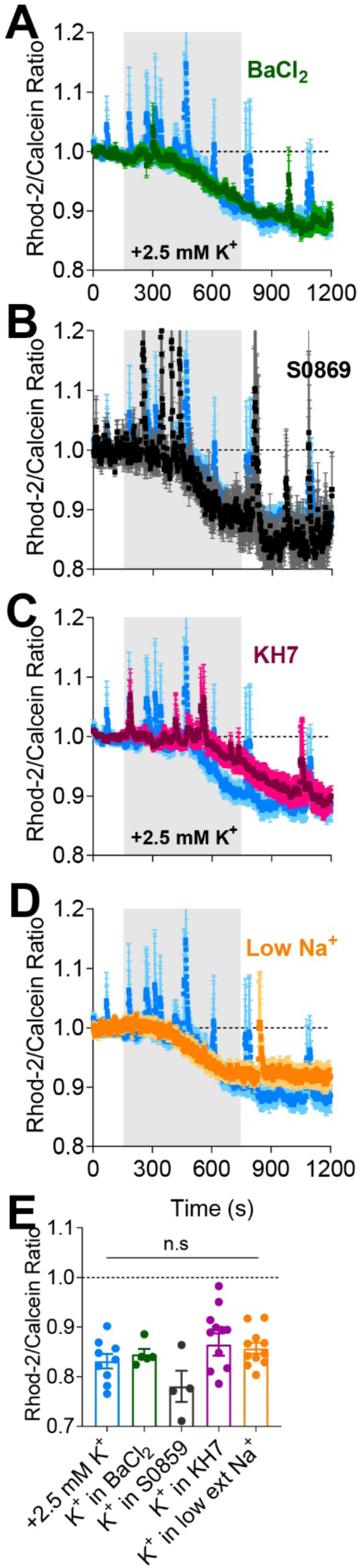
The K^+^ mediated Ca^2+^ decrease does not depend on inward rectifiers, the SCL4A4-sAC pathway or Na^+^ ions. **A-D**) Average summary time series of Rhod-2/Calcein ratio Ca2+ data showing no effect on the K+-mediated decrease when antagonizing inward rectifying potassium channels with BaCl2 (100 μM)*(A)*, blocking SLCaA4 with S0589 (100 μM) *(B)*, antagonizing soluble adenylyl cyclase using KH7 (30 μM) *(C)*, or lowering external Na^+^ to 26.25mM *(D)*.**E**) Summary data of the maximal decrease in astrocyte Ca^2+^ in each experiment. Data are mean +/-SEM, unpaired two-tailed t-tests to the control condition.

